# Connexin Hemichannels with Prostaglandin Release in Anabolic Function of Bone to Mechanical Loading

**DOI:** 10.1101/2021.10.15.464551

**Authors:** Dezhi Zhao, Manuel A. Riquelme, Teja Guda, Chao Tu, Huiyun Xu, Sumin Gu, Jean X. Jiang

## Abstract

Mechanical stimulation, such as physical exercise, is essential for bone formation and health. Here, we demonstrate the critical role of osteocytic Cx43 hemichannels in anabolic function of bone in response to mechanical loading. Two transgenic mouse models, R76W and Δ130-136, expressing dominant-negative Cx43 mutants in osteocytes were adopted. Mechanical loading of tibial bone increased cortical bone mass and mechanical properties in wild-type and gap junction-impaired R76W mice through increased PGE_2_, endosteal osteoblast activity, and decreased sclerostin. These anabolic responses were impeded in gap junction/hemichannel-impaired Δ130-136 mice and accompanied by increased endosteal osteoclast activity. Specific inhibition of Cx43 hemichannels by Cx43(M1) antibody suppressed PGE_2_ secretion and impeded loading-induced endosteal osteoblast activity, bone formation and anabolic gene expression. PGE_2_ administration rescued the osteogenic response to mechanical loading impeded by impaired hemichannels. Together, osteocytic Cx43 hemichannels could be a potential new therapeutic target for treating bone loss and osteoporosis.

## Introduction

Bone as a mechanosensitive tissue is adaptive to mechanical stimuli, which are essential for bone homeostasis, formation, and remodeling (Bonewald, 2011). Reduced mechanical stimulation leads to bone loss and elevated risk of fracture (Lang et al., 2004), while enhanced mechanical stimulation, such as physical exercise, has positive, anabolic impacts on bone tissue, even following a prolonged cessation of stimulation (Erlandson et al., 2012; Warden et al., 2007). The osteocytes embedded in the bone mineral matrix comprise over 90–95% of all bone cells and are thought to be a major mechanoreceptor in the adult skeleton (Bonewald, 2011). Osteocytes detect the mechanical loading-induced alterations of the bone matrix microenvironment and translate them into biological responses to regulate osteoblast and osteoclast activity on the bone surface (Bonewald, 2011; Bonewald and Johnson, 2008).

Connexin (Cx)-forming gap junctions and hemichannels permit small molecules (≤1 kDa) to pass through the cellular membrane, such as prostaglandin E_2_ (PGE_2_) and ATP (Loiselle et al., 2012). Among Cx family members, Cx43 is the predominant Cx subtype expressed in osteocytes (Civitelli, 2008). Cx43 gap junctions allow cell-cell communication between osteocytes or between osteocytes and other bone cell types (Ishihara et al., 2008), and mechanical stimuli increase communication between two adjacent cells through gap junctions (Alford et al., 2003; Cheng et al., 2001). However, osteocytic Cx43 gap junctions are only active at the tips of osteocyte dendritic processes and remain open even without mechanical stimulation (Cusato et al., 2006). In contrast, Cx43 hemichannels, which mediate the communication between the intracellular and the extracellular microenvironment, are highly responsive to mechanical stimulation in osteocytes (Cherian et al., 2005; Jiang and Cherian, 2003). Our previous studies have shown that *in vitro* mechanical stimulation, through fluid flow shear stress (FFSS), increases cell surface expression of Cx43 hemichannels (Cherian et al., 2005; Jiang and Cherian, 2003; Siller-Jackson et al., 2008), and opens Cx43 hemichannels, leading to the release of anabolic factor, PGE_2_ in osteocytes (Cherian et al., 2005; Siller-Jackson et al., 2008). Activation of integrins and PI3K-Akt signaling by FFSS plays an essential role in activating Cx43 hemichannels (Batra et al., 2012; Batra et al., 2014). PGE_2_ released by Cx43 hemichannels acts in an autocrine/paracrine manner to promote gap junction communication through transcriptional regulation of Cx43 (Xia et al., 2010) and blocks glucocorticoid-induced osteocyte apoptosis (Kitase et al., 2010). The opening of Cx43 hemichannels by FFSS also triggers the release of ATP by a protein kinase C-mediated pathway in osteocytes (Genetos et al., 2007). Extracellular PGE_2_ accumulation caused by continuous FFSS exerts a negative feedback, leading to hemichannel closure (Riquelme et al., 2015). However, the biological role of osteocytic Cx43 hemichannels in the anabolic function of mechanical loading has remained largely elusive.

Several bone cell type-specific Cx43 conditional knockout (cKO) mouse models have been reported. Deletion of Cx43 from osteoblasts and osteocytes driven by the Col-2.3-kb α1(I) collagen promoter (Col-2.3kb-Cre; Cx43^−/flx^) attenuated tibial endosteal response to non-physiological mechanical loading, induced by four-point (Grimston et al., 2006) or three-point tibial bending (Grimston et al., 2008). However, deletion of Cx43 in osteochondroprogenitors driven by the Dermo1 collagen promoter (Dermo1-Cre; Cx43^−/flx^)(Grimston et al., 2012) or in osteoblasts/osteocytes driven by the osteocalcin promoter (OCN-Cre; Cx43^flx/flx^)(Zhang et al., 2011) showed an enhanced tibial periosteal response to tibial axial compression (Grimston et al., 2012) or tibial cantilever bending (Zhang et al., 2011). Similarly, deletion of Cx43 in osteocytes driven by an 8-kb dentin matrix protein 1 (DMP1) promoter (DMP1-8kb-Cre; Cx43^flx/flx^) showed enhanced β-catenin levels and correspondingly increased periosteal response to ulna compression (Bivi et al., 2013). Interestingly, endosteal bone formation decreased more in Dermo1-Cre; Cx43^−/flx^ mice (Grimston et al., 2012), but did not change in DMP1-8kb-Cre; Cx43^flx/flx^ mice (Bivi et al., 2013) during mechanical loading. Together, these findings suggest that Cx43 plays a distinct role in the adaptive response to bone loading. However, since Cx43 forms both gap junctions and hemichannels, it has remained largely elusive whether the responses in knockout models could be attributed to either or both types of Cx43-forming channels. Here we dissect the distinctive roles of Cx43 gap junctions and hemichannels using two transgenic mouse models that overexpress dominant-negative Cx43 mutants primarily in osteocytes with the 10 kb DMP1 promoter. The R76W transgenic mouse inhibits gap junctions with enhanced hemichannel function, whereas, Δ130–136 inhibits both gap junctions and hemichannels (Xu et al., 2015). To further delineate the role of osteocytic Cx43 hemichannels under mechanical loading *in vivo*, a monoclonal Cx43(M1) antibody that specifically blocks Cx43 hemichannels was developed. In this study, we unveil a novel role of Cx43 hemichannels in osteocytes and their release of PGE_2_ in mediating anabolic function of the bone in response to mechanical loading.

## Results

### Impairment of Cx43 hemichannels attenuate anabolic responses of tibial bone to mechanical loading

In this study, we used two transgenic mouse models to distinguish the roles of osteocytic Cx43-gap junction channels and hemichannels in osteocytes in bone response to mechanical loading. We injected EB dye into mouse tail veins to determine the activity of Cx43 hemichannels in WT and transgenic mice in response to axial tibial loading. Bone tissue sections around the tibial midshaft region showed that tibial loading increased EB dye uptake in the osteocytes of WT and R76W mice, but not in the osteocytes of Δ130-136 mice (**Figure 1-figure supplement 2A, B**).

We subjected WT and transgenic mice with similar body weights (**Figure 1-figure supplement 3A**) to a 2-week cyclic tibial loading regime. μCT analyses of tibial metaphyseal trabecular bone showed that loading increased bone volume fractions (BV/TV) in WT and R76W mice (**Figure 1A**). In contrast, compared to contralateral, unloaded controls, tibial loading of Δ130-136 mice exhibited a significant reduction of trabecular number (Tb.N) and bone mineral density (BMD), as well as increased trabecular separation (Tb.Sp) during mechanical loading (**Figure 1B, C, E**). However, compared to contralateral, unloaded tibias, trabecular thickness (Tb.Th) was increased in loaded tibias of WT and two transgenic mice (**Figure 1D**). There was no change of structural model index (SMI), indicating that loading did not affect the shape of trabecular bone (**Figure 1F**). Representative μCT images of trabecular bone are shown in **Figure 1G**.

**Figure 1.**
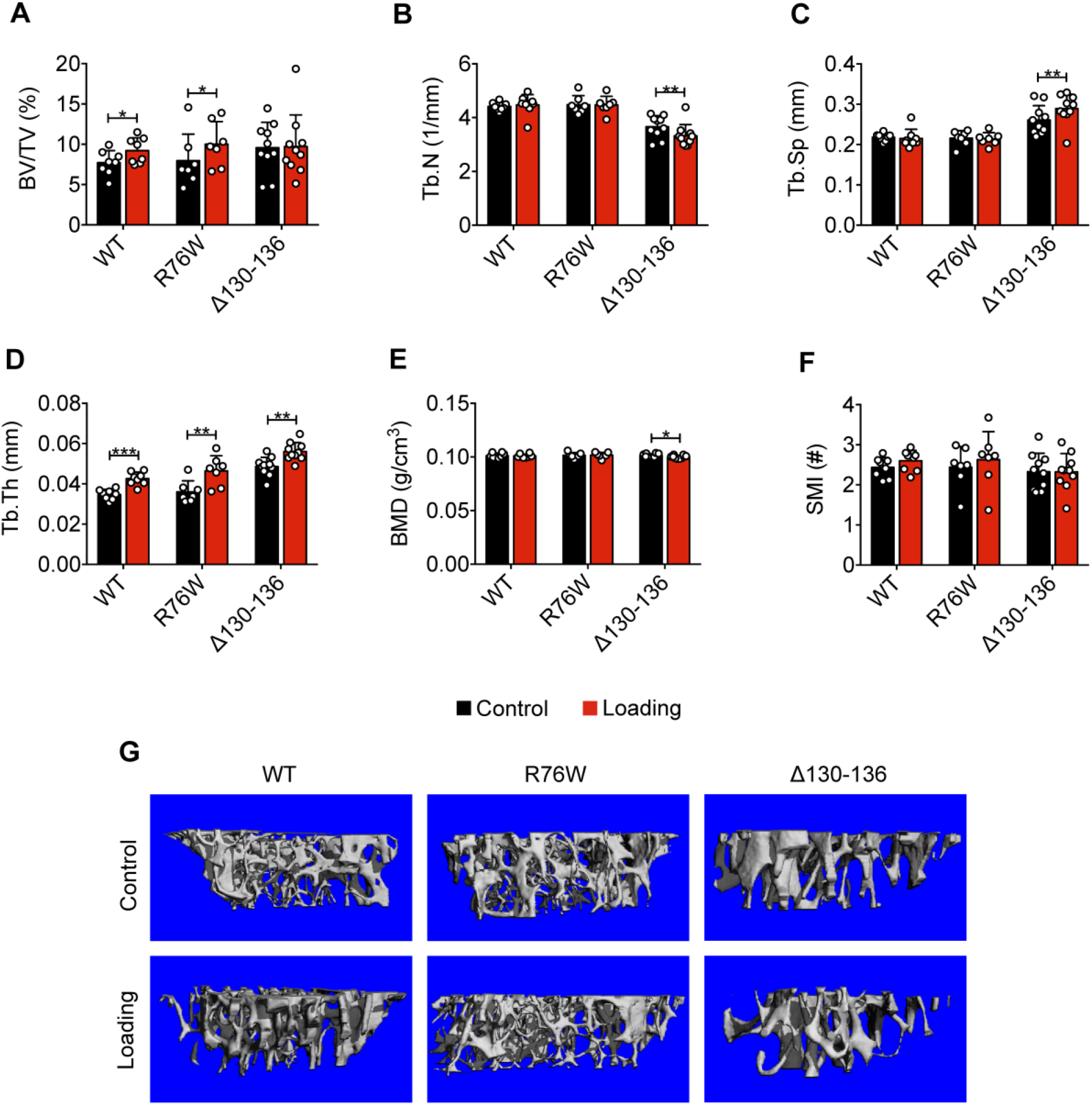
Attenuation or reversal of anabolic responses to mechanical loading in tibial metaphyseal trabecular bone of Δ130-136 mice. μCT was used to assess metaphyseal trabecular bone of WT, R76W, and Δ130-136 mice; **(A)** bone volume fraction, **(B)** trabecular number, **(C)** trabecular separation, **(D)** trabecular thickness, **(E)** bone mineral density, and **(F)** structure model index. n=7-10/group. **(G)** Representative 3D models of the metaphyseal trabecular bone for all groups. Data are expressed as mean ± SD. *, P<0.05; **, P<0.01; ***, P<0.001. Statistical analysis was performed using paired t-test for loaded and contralateral, unloaded tibias.

Similar attenuation of anabolic responses to tibial loading was also observed in cortical bone. μCT analysis was conducted at the midshaft cortical bone (50% site). Loading increased bone area (B.Ar), bone area fraction (B.Ar/T.Ar), and cortical thickness (Ct.Th) in WT and R76W mice (**Figure 2B, C, E**). Although T.Ar was increased by mechanical loading in Δ130-136 (**Figure 2A**), enlarged bone marrow area (M.Ar) (**Figure 2D**) attenuated the ratio of B.Ar/T.Ar (**Figure 2C**). The increased Ct.Th. due to tibial loading was not observed in Δ130-136 mice (**Figure 2E**). Interestingly, the loading caused a decrease of BMD in R76W mice (**Figure 2F**). Torsional strength, predicted by polar moment of inertia (pMOI), was increased as a result of mechanical loading in WT and R76W, but not in Δ130-136 mice (**Figure 2G**). Representative images of cortical bone are shown in **Figure 2H**. Together these data suggested that osteocytic Cx43 hemichannels, not gap junctions, play an important role in anabolic responses of both trabecular and cortical bones to mechanical loading.

**Figure 2.**
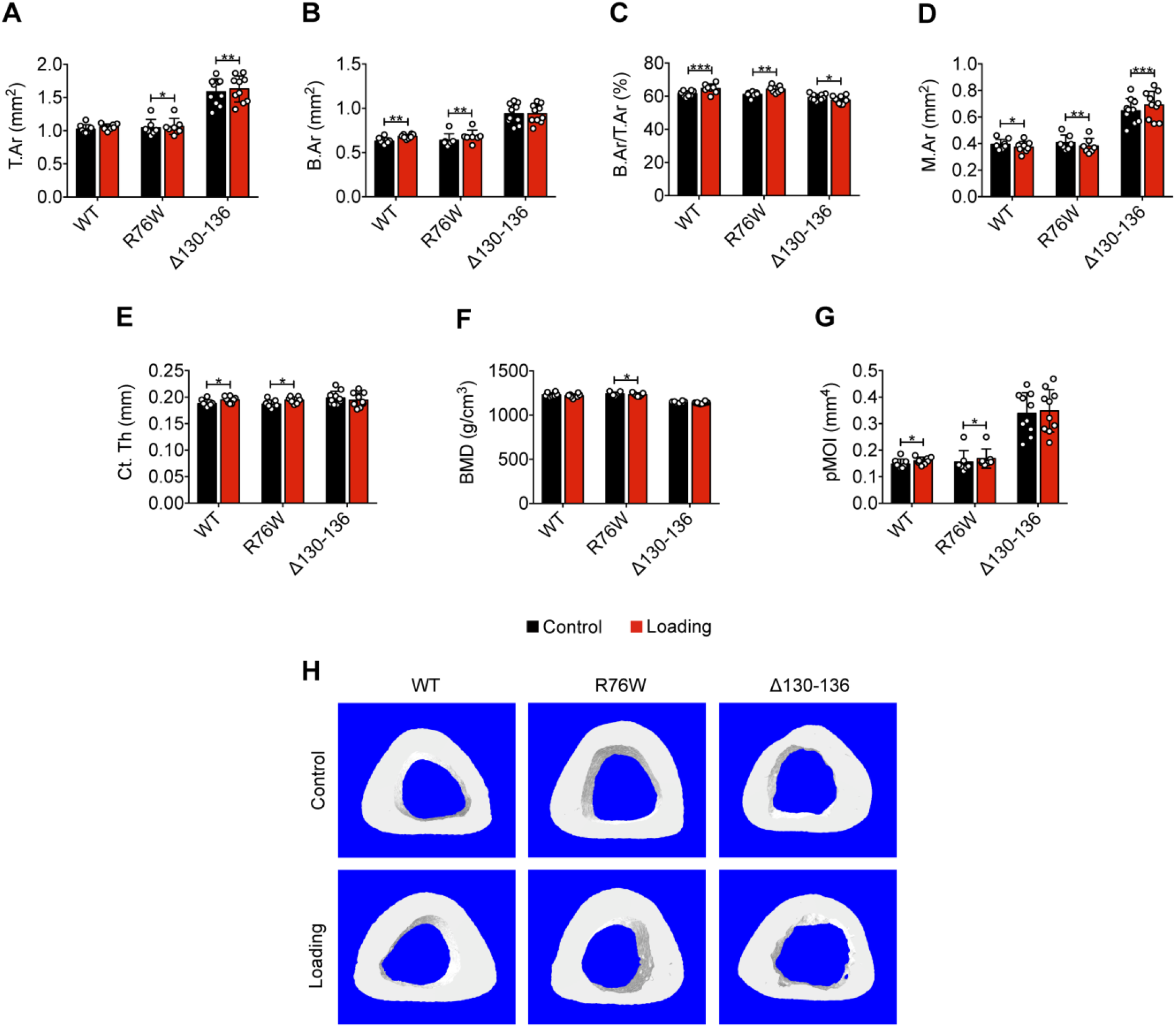
Attenuation or reversal of anabolic responses to mechanical loading in midshaft cortical bone of Δ130-136 mice. μCT was used to assess tibial midshaft cortical bone (50% site) of WT, R76W, and Δ130-136 mice; **(A)** total area, **(B)** bone area, **(C)** bone area fraction, **(D)** bone marrow area, **(E)** cortical thickness, **(F)** bone mineral density, and **(G)** polar moment of inertia. n=7-10/group. **(H)** Representative 3D models of the tibial midshaft cortical bone for all groups. Data are expressed as mean ± SD. *, P<0.05; **, P<0.01; **, P<0.001. Statistical analysis was performed using paired t-test for loaded and contralateral, unloaded tibias.

### Cx43 hemichannels mediate endosteal osteogenic responses to mechanical loading

Dynamic histomorphometric analyses were performed to evaluate periosteal and endosteal bone formation in response to tibial loading. Loading caused a significant increase of endosteal MAR, MS/BS, and BRF/BS compared to contralateral tibias in WT and R76W, but such an increase was not observed in Δ130-136 (**Figure 3A-D**). The decreased endosteal bone formation may partially account for the enlarged bone marrow. Contrary to the endosteal surface, Δ130-136 mice showed a statistically significant osteogenic response on the periosteal surface compared to WT and R76W mice, manifesting a threefold increase in MAR, MS/BS, and BRF/BS during loading (**Figure 3E-H**). Together, these data suggested that impaired osteocytic Cx43 hemichannels in Δ130-136 mice attenuated endosteal bone formation, and enhanced periosteal bone formation upon mechanical loading.

**Figure 3.**
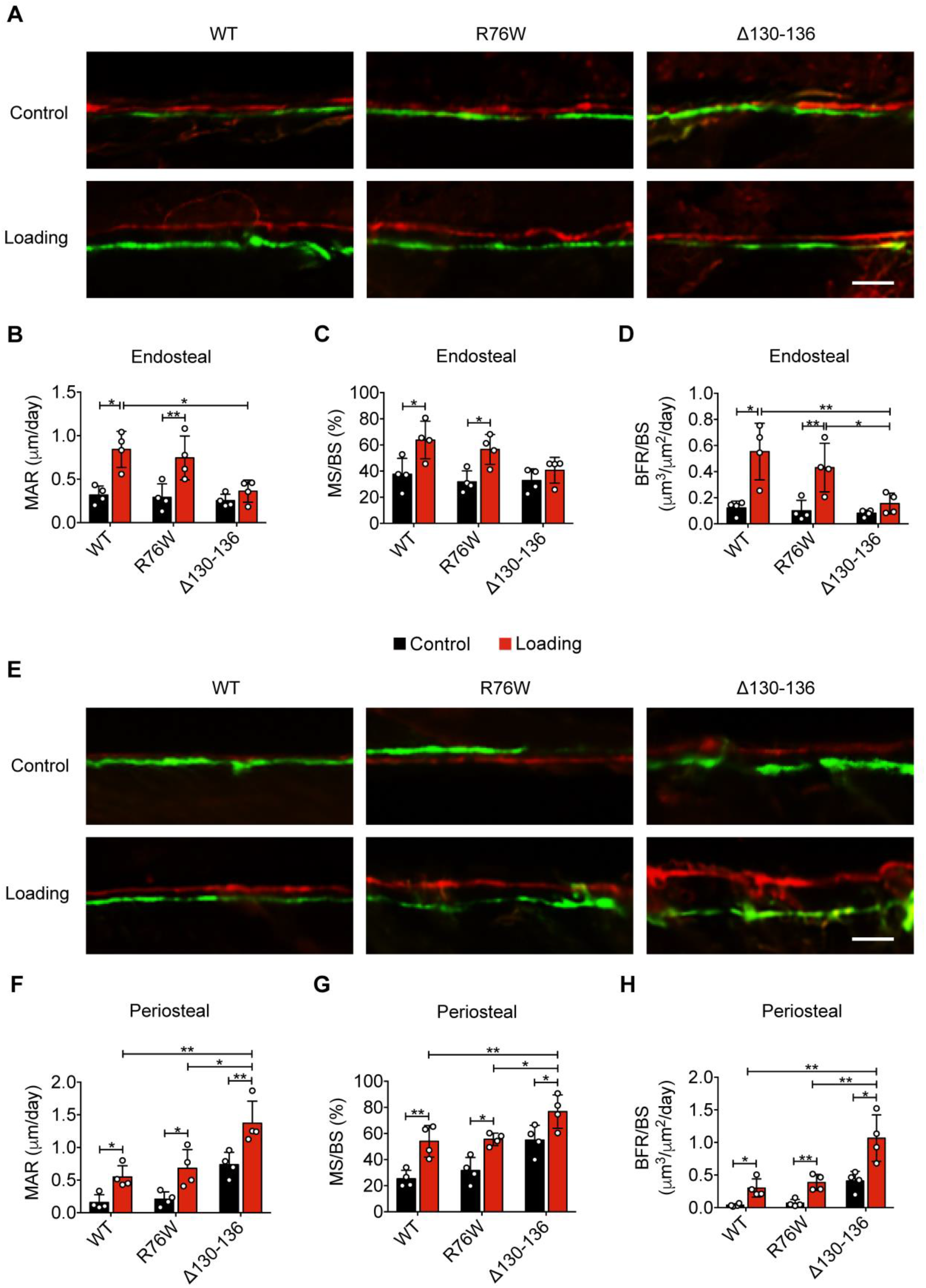
Reduced midshaft endosteal osteogenic responses to mechanical loading in Δ130-136 mice. Dynamic histomorphometric analyses were performed on the tibial midshaft cortical endosteal and periosteal surfaces after 2 weeks of tibial loading of WT, R76W, and Δ130-136 mice. Representative images of calcein (green) alizarin (red) double labeling on **(A)** endosteal and **(E)** periosteal surface. Scale bar: 50 μm. Mineral apposition rate (MAR) **(B and F),** mineralizing surface/bone surface (MS/BS) **(C and G)**, and bone formation rate (BFR/BS) **(D and H)** were assessed for endosteal **(B-D)** and periosteal **(F-H)** surfaces n=4/group. Data are expressed as mean ± SD. *, P<0.05; **, P<0.01. Statistical analysis was performed using paired t-test for loaded and contralateral, unloaded tibias or one-way ANOVA with Tukey test for loaded tibias among different genotypes.

### Impaired Cx43 hemichannels inhibit the loading-induced PGE_2_ secretion and osteoblast activity, and promote osteoclast activity

Cx43 hemichannels mediate PGE_2_ release from osteocytes induced by FFSS *in vitro* (Cherian et al., 2005), and extracellular PGE_2_ is reported to play a key role in the anabolic response to mechanical loading of bone tissue (Jee et al., 1985; Thorsen et al., 1996). We measured PGE_2_ levels in the tibial bone diaphysis and found PGE_2_ levels in mechanically loaded tibias were significantly increased in WT and R76W mice compared to those in contralateral, non-loaded tibias (**Figure 4A**). However, loading had minimal effect on PGE_2_ levels in Δ130-136 mice. Immunohistochemical staining showed that the expression of cyclooxygenase-2 (COX-2), a key enzyme that catalyzes the conversion of arachidonic acid to prostaglandins, was significantly increased in the osteocytes of loaded tibias of WT and R76W mice compared to contralateral, unloaded tibias (**Figure 4B, C** **and Figure 4-figure supplement 1A**). However, the increase of COX-2 in loaded tibias was not observed in Δ130-136 mice. Sost-positive osteocytes decreased significantly in WT and R76W in response to tibial loading, while such decrease in loaded tibias was absent in Δ130-136 mice (**Figure 4D, E** **and Figure 4-figure supplement 1B**). A similar reduction at the mRNA level of the bone was also found in WT and R76W, but absent in Δ130-136 mice (**Figure 4F**). Since Sost, a Wnt receptor antagonist, is a potent inhibitor of osteoblastic activity, we examined osteoblasts on the endosteal surface. WT and R76W mice exhibited an increase of osteoblast numbers on the endosteal surface; in contrast, this increase was absent in Δ130-136 mice (**Figure 4G, H**). Moreover, the levels of mRNA of osteoblastic markers Runx2 and Bglap were higher in the bone of loaded WT and R76W than loaded Δ130-136 mice (**Figure 4I, J**). The mRNA expression of the osteocytic marker Dmp1 in the bone of Δ130-136 mice showed a similar trend of reduction compared to that of control and R76W mice (**Figure 4K**). Contrary to osteoblasts, osteoclast number on the endosteal bone surface was significantly increased in loaded tibias in Δ130-136 mice (**Figure 4L-N**). The data suggested that osteocytic Cx43 hemichannels influence PGE_2_ secretion, key bone marker expression, and osteoblastic and osteoclastic activities on endosteal surfaces in response to mechanical loading.

**Figure 4.**
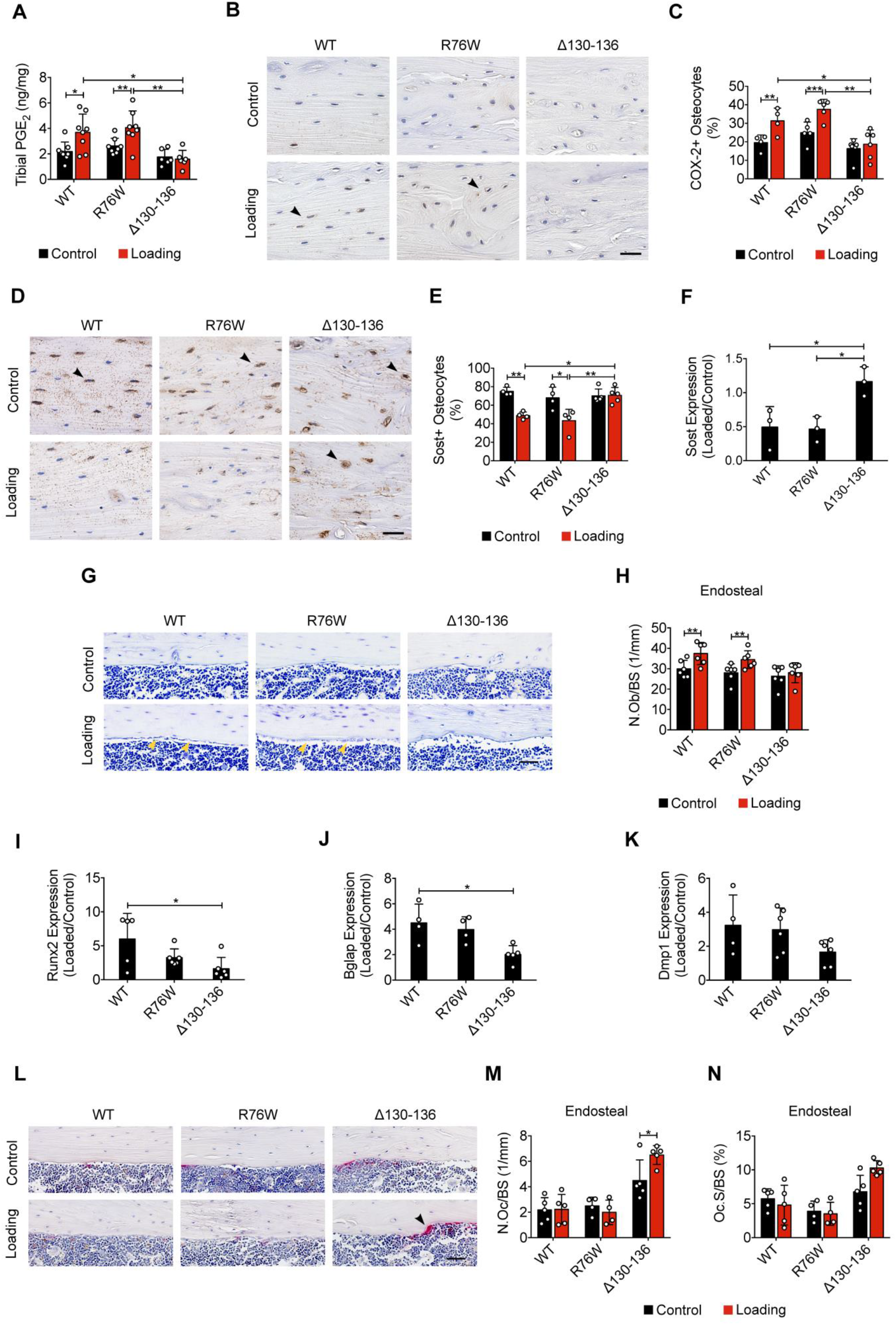
Inhibition of the loading-induced PGE_2_ secretion and osteoblast activity, and promotion of osteoclast activity in Δ130-136 mice. **(A)** ELISA analysis of PGE_2_ in bone marrow-flushed tibial diaphysis after 5 days of tibial loading, in WT, R76W, and Δ130-136 mice. n=6-8/group. **(B and C)** Representative images and quantitative analysis of COX-2-postive osteocytes (black arrows) in the tibial midshaft cortical bone after 2 weeks of loading in WT, R76W, and Δ130-136 mice. Scale bar: 30 μm. n=4-6/group. **(D and E)** Representative images and quantitative analysis of the Sost-positive osteocytes (black arrows) in tibial midshaft cortical bone after 2 weeks of tibial loading in WT, R76W, and Δ130-136 mice. Scale bar: 30 μm. n=4-5/group. **(F)** Gene expression of Sost in bone marrow-flushed tibial diaphysis of WT, R76W, and Δ130-136 mice. n=3/group. **(G and H)** Toluidine blue staining was used to determine the number of endosteal osteoblasts (yellow arrows) on tibial midshaft cortical bone in WT, R76W, and Δ130-136 mice after 2 weeks of loading. Scale bar: 30 μm; n=6/group. mRNA expression of osteoblast markers, Runx2 **(I)** and Bglap (**J**), and osteocyte marker, Dmp1 **(K**) in bone marrow-flushed tibial diaphysis of WT, R76W, and Δ130-136 mice. n=4-6/group. **(L)** Representative images of tibial midshaft endosteal surface stained for TRAP (black arrows). Scale bar: 30 μm. (**M and N**) Histomorphometric quantitation of osteoclasts per bone perimeter **(M)** and osteoclast surface per bone perimeter **(N)** (n=4-5/group). All quantitative data are expressed as mean ± SD. *, P<0.05; **, P<0.01; ***, P<0.001. Statistical analysis was performed using paired t-test for loaded and contralateral tibias or one-way ANOVA with Tukey test for loaded tibias among different genotypes.

### Cx43 hemichannel-blocking antibody impairs the anabolic effects of mechanical loading on trabecular and cortical bones

We have developed a polyclonal antibody, Cx43(E2), that targets an extracellular loop domain of Cx43 and specifically blocks osteocytic Cx43 hemichannels (Siller-Jackson et al., 2008). We recently developed a specific mouse monoclonal blocking antibody Cx43(M1) to investigate the roles of hemichannels *in vivo*. Gap junction channels and hemichannels were assayed using dye coupling (**Figure 5-figure supplement 1A**) and dye uptake assays (**Figure 5-figure supplement 1B**), respectively. Both Cx43(E2) and Cx43(M1) had minimal effects on gap junction channels as indicated by comparable levels of dye transfer with red-orange AM dye in MLO-Y4 cells (**Figure 5-figure supplement 1A**). Conversely, FFSS-induced hemichannel opening, as determined by EtBr uptake, was inhibited by Cx43(E2) and Cx43(M1) antibodies in MLO-Y4 cells (**Figure 5-figure supplement 1B**). The extent of inhibition is comparable between Cx43(E1) and Cx43(M1) antibodies. To ensure antibody delivery to osteocytes, we labeled tibial bone sections with a rhodamine-conjugated anti-mouse secondary antibody. Strong antibody signals were primarily detected in osteocytes in cortical bone for Cx43(M1)-injected mice, but not in IgG-injected ones (**Figure 5-figure supplement 1C**). Interestingly, low levels of Cx43(M1) were detected in trabecular bone (**Figure 5-figure supplement 1D**). The hemichannel opening detected by EB fluorescence was found only in loaded tibial bone, and this uptake was almost completely blocked by the Cx43(M1) antibody (**Figure 5-figure supplement 1E, F**). Hence, these studies established the feasibility of using Cx43(M1) antibody to assess the role of Cx43 hemichannels *in vivo* using the tibial loading model.

WT mice with similar body weight (**Figure 5-figure supplement 2A**) were randomly allocated to Cx43(M1) or vehicle groups. A slight decline in body weight was found during the first week of the study, but was stabilized by the second week (**Figure 5-figure supplement 2A**). The antibody had a negligible effect on body weight (**Figure 5-figure supplement 2A**). μCT analysis of tibial metaphyseal trabecular bone showed that Cx43(M1) treatment significantly abated the loading-induced increase of Tb.N and decrease of Tb.Sp (**Figure 5B, C**) as compared to the vehicle-treated group, while Cx43(M1) did not manifest significant differences in the Tb.Th, BV/TV, and BMD, in response to loading (**Figure 5D-F**). There was no change of SMI in both Cx43(M1) or vehicle groups (**Figure 5G**). Representative images of trabecular bone are shown in **Figure 5A**.

**Figure 5.**
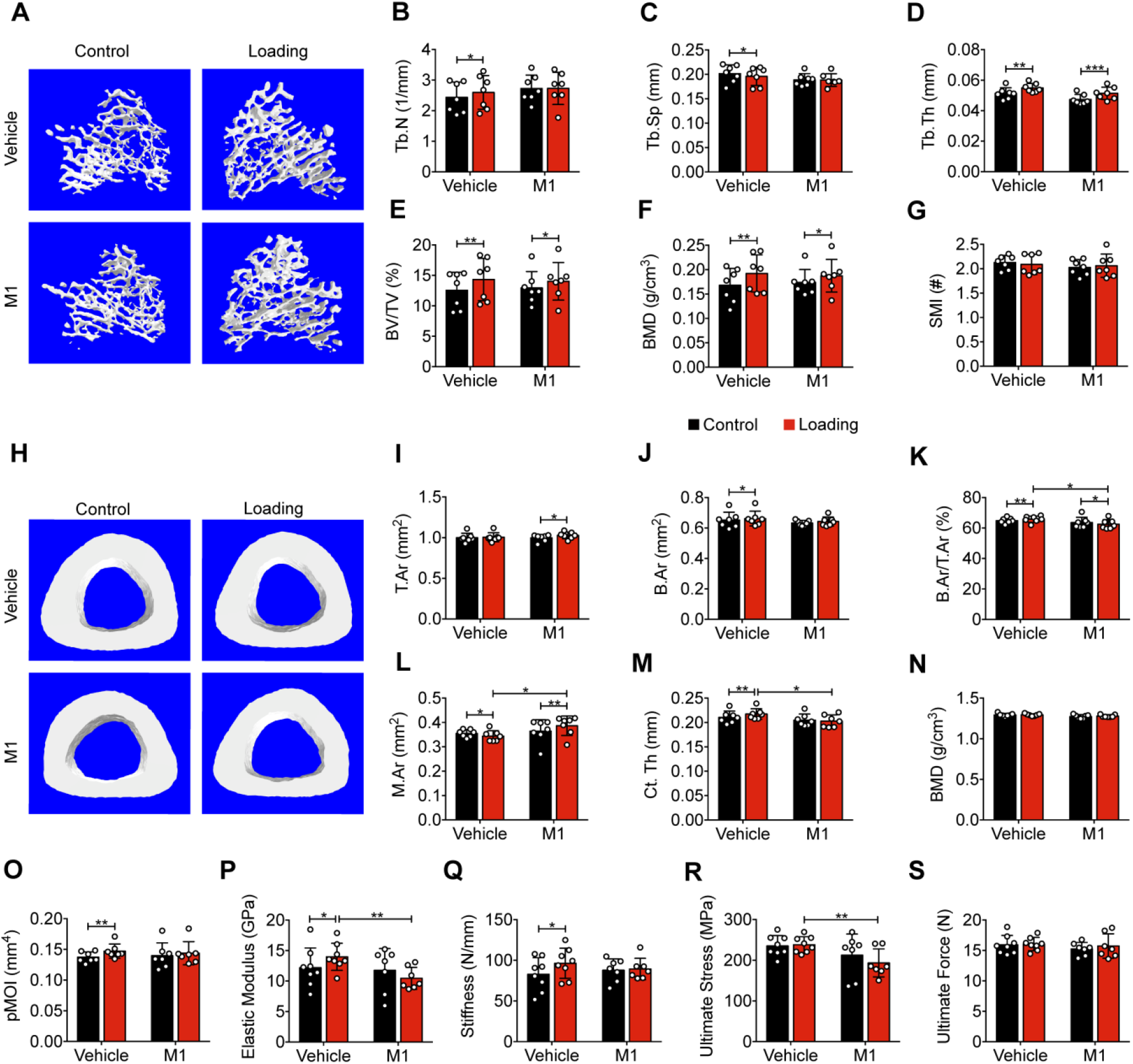
Inhibition of Cx43 hemichannels by Cx43(M1) antibody impairs anabolic effects of mechanical loading on trabecular and cortical bones. **(A)** Representative 3D models of the metaphyseal trabecular bone of vehicle and Cx43(M1)-treated mice. **(B-G)** μCT was used to assess structural parameters of trabecular bone; **(B)** trabecular number, **(C)** trabecular separation, **(D)** trabecular thickness, **(E)** bone volume fraction, **(F)** bone mineral density, and **(G)** structure model index in vehicle and Cx43 (M1)-treated mice. n=7/group. **(H)** Representative 3D models of the tibial midshaft cortical bone (50% site) in vehicle and Cx43(M1)-treated mice. **(I-N)** μCT was used to assess structural parameters of cortical bone; **(I)** total area, **(J)** bone area, **(K)** bone area fraction, **(L)** bone marrow area, **(M)** cortical thickness, **(N)** bone mineral density and **(O)** polar moment of inertia in vehicle and Cx43(M1)-treated mice. n=7/group. **(P-S)** The three-point bending assay was performed for tibial bone of vehicle and Cx43 (M1)-treated mice; **(P)** elastic modulus, **(Q)** stiffness, **(R)** ultimate stress, and **(S)** ultimate force. n=7-8/group. Data are expressed as mean ± SD. *, P<0.05; **, P<0.01; ***, P<0.001. Statistical analysis was performed using paired t-test for loaded and contralateral or unpaired t-test for loaded tibias between vehicle- and Cx43(M1)-treated groups.

Cx43(M1) treatment attenuated the anabolic response to mechanical loading in midshaft cortical bone. The increase of B.Ar, B.Ar/T.Ar and Ct.Th by tibial loading was attenuated in the Cx43(M1)-treated group (**Figure 5J, K, M**), and consequently, the increase of pMOI was also attenuated in this group (**Figure 5O**). Similar to Δ130-136 mice, larger M.Ar and lower B.Ar/T.Ar ratios were found in loaded tibias of the Cx43(M1)-treated group when compared to the vehicle group (**Figure 5K, L**). Although Cx43(M1) further increased T.Ar by mechanical loading (**Figure 5I**), enlarged M.Ar (**Figure 5L**) significantly reduced the ratio of B.Ar/T.Ar (**Figure 5K**). Thus, Ct.Th did not increase and was even lower in loaded tibia compared to vehicle loaded tibias (**Figure 5M**). However, BMD was not changed by mechanical loading (**Figure 5N**). Three-point bending analyses revealed a significant increase of elastic modulus and stiffness only in the vehicle group (**Figure 5P, Q**). For loaded tibias, the elastic modulus and ultimate stress in the vehicle group were greater than the Cx43(M1) group (**Figure 5P, R**). Mechanical loading did not change ultimate force in either vehicle or Cx43(M1)-treated group (**Figure 5S**). Representative images of cortical bone are shown in **Figure 5H**. These results are consistent with those obtained from Δ130-136 mice, suggesting the critical roles of Cx43 hemichannels in the anabolic effects of mechanical loading of cortical bone.

### Blocking Cx43 hemichannels by Cx43(M1) inhibits the load-induced increase in midshaft endosteal osteogenesis

Bone formation in response to tibial loading was evaluated in vehicle and Cx43(M1)-treated mice. The vehicle group exhibited increased endosteal MAR, MS/BS, and BRF/BS compared to contralateral, unloaded controls, whereas this response was absent in the Cx43(M1) group (**Figure 6A-D**). In contrast, loading increased bone formation in vehicle and Cx43(M1)-treated groups on the periosteal surface (**Figure 6E-H**). Cx43(M1) treatment induced a greater increase in periosteal MS/BS (**Figure 6G**). The results showed that impaired Cx43 hemichannels attenuated endosteal bone formation, but enhanced periosteal bone formation, induced by mechanical loading.

**Figure 6.**
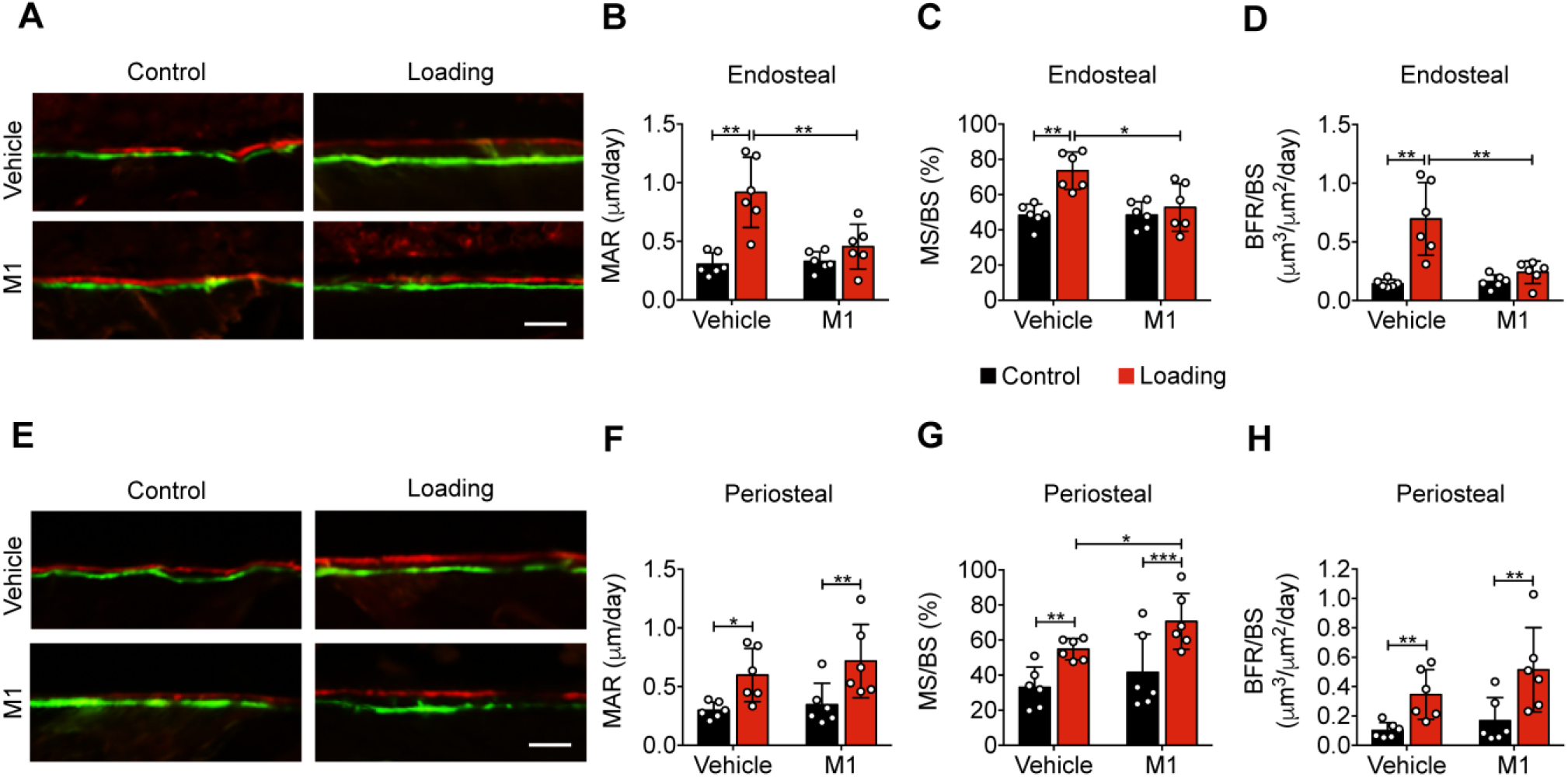
Cx43(M1) inhibits the load-induced increase in midshaft endosteal osteogenesis. Dynamic histomorphometric analyses were performed on the tibial midshaft cortical endosteal **(A-D)** and periosteal **(E-H)** surfaces after 2 weeks of loading in vehicle and Cx43(M1)-treated mice. **(A and E)** Representative images of calcein (green) alizarin (red) double labeling on **(a)** endosteal and **(E)** periosteal surface Scale bar: 50 μm. Mineral apposition rate (MAR), mineralizing surface/bone surface (MS/BS), and bone formation rate (BFR/BS) were assessed for **(B-D)** endosteal and **(F-H)** periosteal surfaces (n=6/group). Data are expressed as mean ± SD. *, P<0.05; **, P<0.01; ***, P<0.001. Statistical analysis was performed using paired t-test for loaded and contralateral, unloaded tibias or unpaired t-test for loaded tibias between vehicle and Cx43(M1)-treated groups.

### Blocking Cx43 hemichannels by Cx43(M1) impedes the loading-induced increased PGE_2_ secretion, bone marker expression, and endosteal osteoblastic activity, and decreased osteoclastic activity

Tibial loading significantly increased PGE_2_ expression in tibial bone, and this increase was not observed with Cx43(M1) antibody treatment (**Figure 7A**). Immunohistochemical staining of tibial cortical bone showed a significant increase of COX-2 positive osteocytes by mechanical loading, and such increase was not detected in the Cx43(M1)-treated group (**Figure 7B, C** **and Figure 7-figure supplement 1A**). COX-2 bone mRNA levels detected by RT-qPCR exhibited close to a 5-fold increase due to tibial loading, and this increase was not detected in Cx43(M1) treated bone samples (**Figure 7D**). Loading caused a significant decrease of Sost-positive osteocytes in tibial bone in the vehicle group, and Cx43(M1) antibody abated the load-induced decrease of Sost-positive osteocytes (**Figure 7E, F and Figure 7-figure supplement 1B**). Sost bone mRNA also showed a significant decrease due to loading in the vehicle group compared to the Cx43(M1)-treated group (**Figure 7G**). We next determined another mechanical response protein, β-catenin expression, in osteoblasts. Tibial loading resulted in a robust increase in endosteal β-catenin-positive osteoblasts and tibial β-catenin gene expression in the vehicle group; in contrast, such increase was absent in the Cx43(M1) group (**Figure 7H-J** **and Figure 7-figure supplement 1C**). Consistent with β-catenin expression, endosteal osteoblast number only increased in the vehicle group (**Figure 7K, L**). Moreover, the increase of gene expression of the osteoblastic marker, Bglap, was greater in the bone of the vehicle group compared to the Cx43(M1) group (**Figure 7M**). Interestingly, increased osteoclast activity was also found in the Cx43(M1) group (**Figure 7K, N, O**). The results showed that under mechanical loading, inhibition of Cx43 hemichannels by Cx43(M1) antibody impedes the PGE_2_ release and Sost decrease in osteocytes. This was associated with inhibited β-catenin and Bglap expression and osteoblast activity on the endosteal surface.

**Figure 7.**
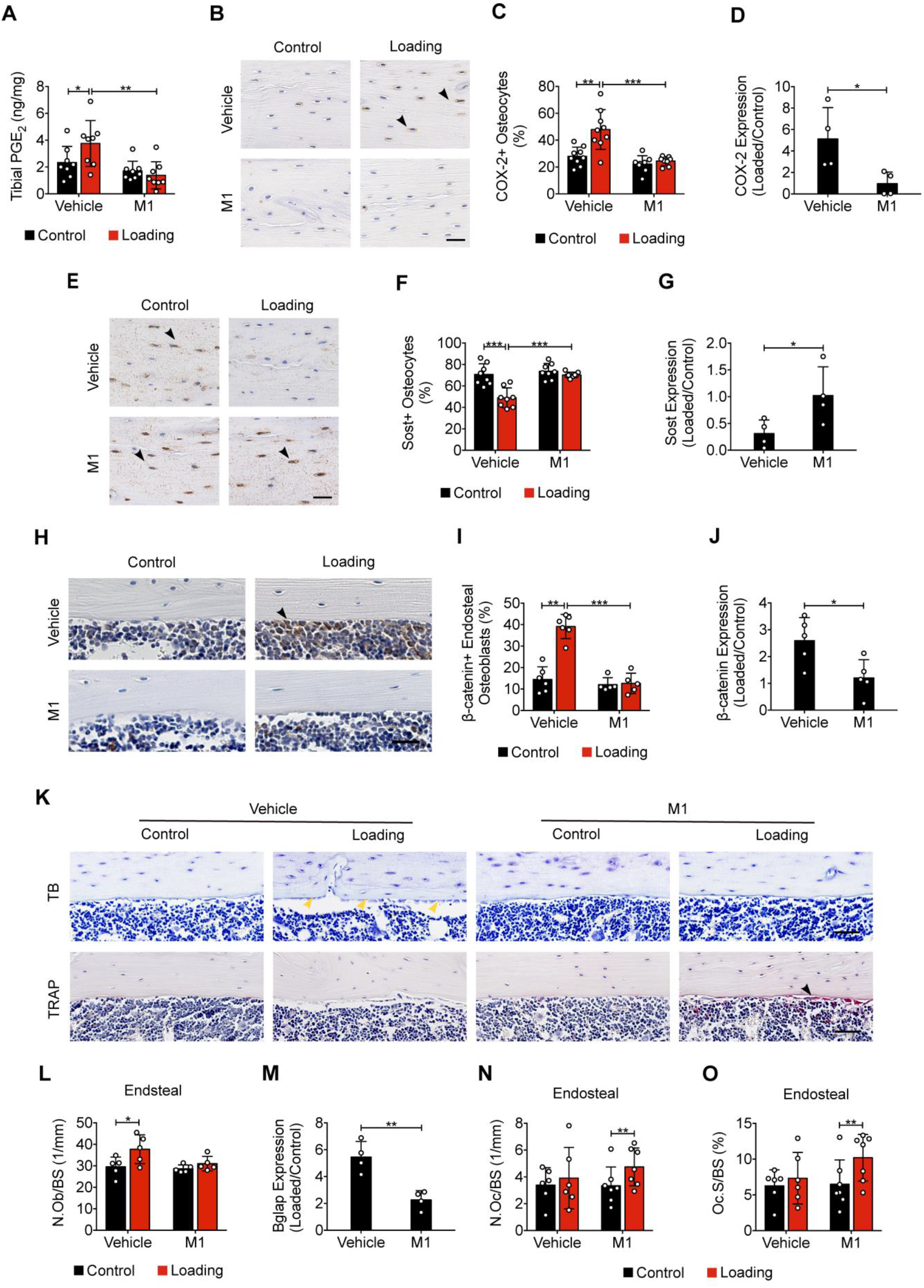
Cx43(M1) impedes the loading-induced increased PGE_2_ secretion and osteoblast activity, and decreased osteoclast activity. **(A)** ELISA analysis of PGE_2_ in bone marrow-flushed tibial diaphysis after 5 days of mechanical loading in vehicle and Cx43(M1)-treated mice (n=8/group). (**B and C**) Representative images and quantitative analysis of COX-2-postive osteocytes (yellow arrows) in tibial midshaft cortical bone after 2 weeks of loading in vehicle and Cx43(M1)-treated mice. Scale bar: 30 μm. n=8-9/group. **(D)** COX-2 mRNA determined by RT-qPCR in bone marrow-flushed tibial diaphysis of vehicle and Cx43(M1)-treated mice. n=4/group. **(E-F)** Representative images and quantitative analysis of the Sost-positive osteocytes (yellow arrows) in tibial midshaft cortical bone after 2 weeks of mechanical loading in vehicle and Cx43(M1)-treated mice. Scale bar: 30 μm (n=8/group). **(G)** Sost mRNA determined by RT-qPCR from bone marrow-flushed tibial diaphysis of vehicle and Cx43(M1)-treated mice. n=4/group. **(H and I**) Representative images and quantitative analysis of the β-catenin positive periosteal cells (black arrows) on tibial midshaft endosteal surface after 2 weeks of loading in vehicle and Cx43(M1)-treated mice. Scale bar: 20 μm; n=5-6/group. **(J)** β-catenin mRNA determined by RT-qPCR in bone marrow-flushed tibial diaphysis of vehicle and Cx43(M1)-treated mice. n=4/group. (**K**) Representative images of tibial midshaft endosteal surface stained for toluidine blue (top panel) or TRAP (low panel). The yellow arrows indicate osteoblasts and the black arrows indicate the TRAP-positive osteoclasts. Scale bar: 30 μm. (**L**) Histomorphometric quantitation of osteoblast per bone perimeter (n=5-7/group). **(M)** Bglap mRNA determined by RT-qPCR in bone marrow-flushed tibial diaphysis of vehicle and Cx43(M1)-treated mice. n=4/group. (**N and O**) Histomorphometric quantitation of osteoclast per bone perimeter **(N)** and osteoclast surface per bone perimeter **(O)** (n=5-7/group). Data are expressed as mean ± SD. *, P<0.05; **, P<0.01; ***, P<0.001. Statistical analysis was performed using paired t-test for loaded and contralateral tibias, unpaired t-test for loaded tibias between vehicle and Cx43(M1)-treated groups.

### PGE_2_ rescues impeded osteogenic responses to mechanical loading by impaired Cx43 hemichannels

Intermittent PGE_2_ treatment has been reported to increase both trabecular and cortical bone mass (Jee et al., 1985; Tian et al., 2007). To explore whether the attenuated anabolic function of bone to mechanical loading with Cx43(M1) is caused by inhibited PGE_2_ released by Cx43 hemichannels, PGE_2_ was IP injected into the vehicle control and Cx43(M1) treated mice. The mice in the control and treated groups had comparable body weights to minimize variations in tibial bone sizes before loading (**Figure 8-figure supplement 1A**). μCT analysis showed that there was no difference in trabecular morphometric parameters among the four groups (**Figure 8-figure supplement 2**). Contrary to trabecular bone, PGE_2_ treatment impeded the significant reduction of B.Ar/T.Ar ratio, decreased M.Ar and increased Ct.Th in Cx43(M1)-treated loaded tibias, although there were no significant differences in T.Ar, B.Ar, and BMD in all four groups (**Figure 8B-G**). Representative images of cortical bone are shown in **Figure 8A**. Interestingly, PGE_2_ did not further enhance anabolic bone responses in control, loaded mice. Together, these results demonstrate that administration of PGE_2_ significantly rescues impeded anabolic responses of cortical bone to mechanical loading as a result of Cx43 hemichannel inhibition.

**Figure 8.**
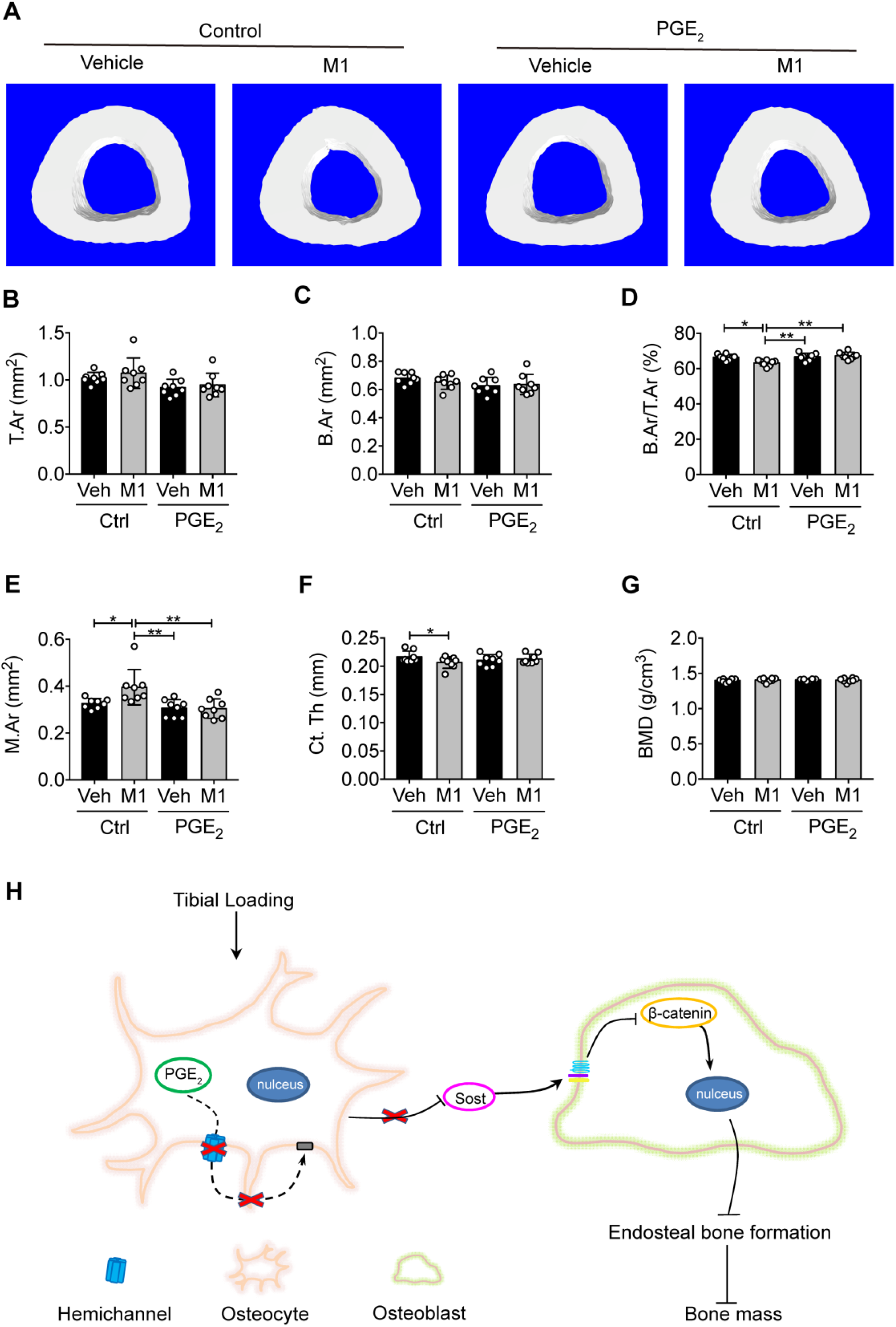
PGE_2_ rescues the osteogenic response to mechanical loading with the impairment of Cx43 hemichannels in cortical bone. **(A)** Representative 3D models of the tibial midshaft cortical bone (50% site) in vehicle and Cx43(M1)-treated mice treated with 1 mg/kg/day PGE_2_ or vehicle control. (**B-G**) μCT was used to assess tibial midshaft cortical bone; **(B)** total area, **(C)** bone area, **(D)** bone area fraction, **(E)** bone marrow area, **(F)** cortical thickness, and **(G)** bone mineral density. n=8/group. Data are expressed as mean ± SD. *, P<0.05; **, P<0.01; ***, P<0.001. Statistical analysis was performed using one-way ANOVA with Tukey test. (**H**) Schematic diagram illustrating the mechanistic roles of osteocytic Cx43 hemichannels in mediating anabolic responses to tibial loading. Briefly, Cx43 hemichannels mediate the release of PGE_2_ by mechanical loading, leading to suppression of Sost expression with enhanced β-catenin expression and osteogenesis on the endosteal surface. The inhibition of Cx43 hemichannels impedes the loading-induced PGE_2_ secretion and anabolic function of mechanical loading on bone tissue.

## Discussion

In this study, we unveiled the distinctive roles of two types of Cx43 channels in responses to mechanical loading using both transgenic mouse models and Cx43 hemichannel-blocking antibodies, and demonstrated the critical role of Cx43 hemichannels in mediating the anabolic, or bone forming, function of bone upon mechanical loading. Moreover, PGE_2_, a factor released by Cx43 hemichannels in response to mechanical stimulation, rescued the impeded anabolic effects on bone by tibial loading as a result of impaired hemichannels.

We determined bone structural and biomechanical properties, new bone formation, and metabolism using an established bone mechanical loading model, axial tibial compression, in two transgenic mouse models expressing Cx43 dominant negative mutants R76W and Δ130-136 in osteocytes. We observed that Δ130-136 mice, with inhibited osteocytic Cx43 hemichannels, attenuated anabolic bone responses to mechanical loading in both trabecular and cortical bones. The attenuation of bone formation and osteoblastic activities primarily occurred on the tibial endosteal surface, while increased bone formation was observed on the periosteal surface. The activities of osteoblasts and osteoclasts on periosteal and endosteal surfaces, respectively, lead to an enlarged bone marrow area and decreased bone to tissue area ratio. Due to impaired osteocytic gap junction channels and hemichannels in Δ130-136 mice, but only impaired gap junction channels in R76W mice, we postulated that osteocytic Cx43 hemichannels, not gap junctions in osteocytes, are likely to play a predominant role in anabolic bone response to mechanical loading.

The role of Cx43 hemichannels was further validated by the Cx43(M1) antibody, a potent monoclonal antibody that effectively inhibits osteocytic Cx43 hemichannels both *in vitro* and *in vivo*. Remarkably, treatment with Cx43(M1) only twice in a span of two weeks significantly attenuated anabolic effects to mechanical loading, with greater effects in cortical bones. Interestingly, Cx43(M1) treatment not only attenuated, but even reversed anabolic effects in cortical bone, similar to Δ130-136 mice. A similar impediment of the rate of bone formation and mineral apposition to tibial loading was observed on the endosteal surface with Cx43(M1) treatment. Previous studies have shown that cortical bone modeling/remodeling is more pronounced at the endosteal surface (Birkhold et al., 2017), and mature bones respond to mechanical loading through changes on endosteal surfaces (Bass et al., 2002). Similar findings were also noted in Cx43 cKO mouse models; mice lacking Cx43 in osteoblasts and osteocytes showed an attenuated increase in endosteal bone formation during four-point or three-point tibial bending (Grimston et al., 2008; Grimston et al., 2006). Mice lacking Cx43 in osteochondroprogenitors showed a greater extent of decrease in endosteal formation during tibial compression loading (Grimston et al., 2012). In our study, notably, endosteal MAR and BFR/BS were not responsive to tibial loading in Δ130-136 mice and the Cx43(M1) group, suggesting not only increased osteoblastic activity, but also that increased osteoblast number was impeded by the impairment of Cx43 hemichannels. Moreover, histomorphometric analysis further confirmed a lack of response in osteoblast number and marker genes in Δ130-136 mice and Cx43(M1)-treated mice. The difference between Cx43 cKO and our transgenic models and Cx43(M1) treatment could be caused by aberrant, compensatory effect of other pathways as a result of Cx43 deletion. Thus, our results suggest that axial compression loading promotes osteoblast recruitment and differentiation on the endosteal surface, an anabolic effect likely mediated by mechanosensitive Cx43 hemichannels.

Interestingly, contrary to our hypothesis, axial load increased periosteal bone formation and total tissue area on the tibial midshaft in Δ130-136 mice. This observation was also reported in cKO mouse models with Cx43 deletion in osteoblasts and osteocytes under tibial axial compression (Grimston et al., 2012) and tibial cantilever bending (Zhang et al., 2011). Similarly, deletion of Cx43 in osteocytes also showed an enhanced periosteal response to ulnar compression (Bivi et al., 2013). The enhanced periosteal bone formation was further observed when hemichannels were inhibited by Cx43(M1). These results posit the role of Cx43 hemichannels in periosteal bone formation during mechanical loading. Our previous study showed that Δ130-136 mice have more periosteal bone apposition than WT and R76W mice, suggesting that periosteal osteoblasts in Δ130-136 mice are more active and sensitive than WT and R76W mice (Xu et al., 2015). Thus, osteocytic Cx43 hemichannels exert differential roles in controlling osteogenic osteoblastic activities on periosteal and bone resorping osteoclastic activity on endosteal surfaces, respectively. The consequence of the disruption of coordinated activities by hemichannel inhibition results in an enlarged bone marrow cavity. This is likely an adaptive response due to compromised cortical bones resulting from impaired hemichannels and consequently lower extracellular PGE_2_. From a mechanical point of view, increased bone marrow area and cortical bone size allow the bone to respond to high stress levels (Sharir et al., 2008). Thus, the role of osteocytic Cx43 hemichannels in regulating the osteogenic response on endosteal surfaces is distinct from its role on periosteal surfaces.

Besides cortical bone, the anabolic response of trabecular bone to tibial loading in Δ130-136 mice was also attenuated or even reversed, as observed in trabecular number, separation and bone density. However, the compromised response of trabecular bone to mechanical loading was less evident in Cx43(M1) treated mice, except for trabecular number and trabecular separation. One possible explanation is that the accessibility and binding of the antibody to osteocytes in trabecular bone may not be as efficient as in cortical bone. Indeed, we could clearly detect Cx43(M1) on the surface of osteocytes in cortical bone. However, the binding in trabecular bone is much weaker than that in cortical bone. Since Haversian canals containing blood vessels provide supply to the osteocytes in cortical, but not in trabecular bone (Dahl and Thompson, 2011), it is plausible that the delivery of Cx43(M1) to the bone is mediated primarily by the Haversian canal system.

Previous studies have reported that *in vitro* Cx43 hemichannel opening induced by FFSS mediates the release of PGE_2_ (Cherian et al., 2005), a critical factor for anabolic function of bone in response to mechanical loading (Jee et al., 1985; Thorsen et al., 1996). On the contrary, PGE_2_ release by FFSS is inhibited by a potent hemichannel-blocking rabbit polyclonal antibody Cx43(E2) (Siller-Jackson et al., 2008). Here, we found that PGE_2_ levels and osteocytic COX-2 expression were increased by tibial loading in WT and R76W mice, but such an increase was not detected in Δ130-136 mice. The use of hemichannel-blocking monoclonal Cx43(M1) antibody further confirmed the role of the hemichannels in the release of PGE_2_ in bone *in situ*. In accordance with our observation, the reduced release of PGE_2_ was reported in calvarial cells isolated from Cx43 cKO mice driven by the Col-2.3-kb α1(I) collagen promoter after mechanical stretching (Grimston et al., 2006). The increase in *Cox-2* gene expression in Cx43 cKO mice driven by the 8-kb DMP1 promoter is attenuated after axial tibial compression (Grimston et al., 2012).

We showed that inhibited release of PGE_2_ in Δ130-136 mice and in the Cx43 (M1) group was accompanied by an attenuated endosteal bone response to mechanical loading. Moreover, PGE_2_ injection rescued the anabolic responses of cortical bone to mechanical loading impeded by Cx43(M1), including the ratio of bone area to tissue area, cortical thickness, and bone marrow area. PGE_2_ is a skeletal anabolic factor, and its synthesis and release are highly responsive to mechanical stimulation in osteocytes (Cherian et al., 2005; Jiang and Cherian, 2003). Using the microdialysis technique, a rapid and significant increase of PGE_2_ levels in the proximal tibial metaphysis was observed in response to dynamic mechanical loading in healthy women (Thorsen et al., 1996). Furthermore, intermittent PGE_2_ treatment increases endosteal bone formation (Jee et al., 1985) and bone mass (Tian et al., 2007). Conversely, inhibition of PGE_2_ by a COX-2 inhibitor blocks endosteal tibial bone formation induced by mechanical loading in rats (Forwood, 1996). Here, we demonstrate that PGE_2_ is indeed involved in the anabolic action of hemichannels in response to mechanical loading. Interestingly, PGE_2_ administration did not provide an additional increase in the cortical bone of mice in loaded vehicle control group. It is likely that extracellular PGE_2_ released by osteocytes by normal exercise (mechanical loading) is sufficient to promote bone formation and additional extracellular PGE_2_ would not further increase cortical bone mass.

Increased PGE_2_ by mechanical stimuli is reported to bind to EP4 receptor and reduce Sost expression (Galea et al., 2011). Sost, a Wnt signaling antagonist (Semenov et al., 2005), is a key regulator of mechanotransduction in bone. Sost, secreted primarily by osteocytes, acts upon osteoblasts in a paracrine manner to inhibit bone formation (Poole et al., 2005) through its binding to the Wnt co-receptor Lrp5/6 (Li et al., 2005) and suppressing β-catenin (Sawakami et al., 2006). Sost gene and protein expression is suppressed by mechanical loading, and is accompanied by increased bone formation(Moustafa et al., 2012; Robling et al., 2008). We observed suppressed Sost expression in WT and R76W mice by tibial loading; however, the suppressive effect of Sost disappeared in Δ130-136 mice and the Cx43(M1)-treated mice. Correspondingly, the increased β-catenin expression and osteoblast activity observed in WT and R76W mice was abated in Δ130-136 and the Cx43(M1) mice. These results indicate that PGE_2_ released by Cx43 hemichannels in osteocytes is a likely factor that participates in the bone anabolic response to mechanical stimuli. We previously showed that PGE_2_ released from osteocytes via Cx43 hemichannels exerts autocrine effects via the EP2/4 receptor during mechanical stimulation (Xia et al., 2010). This study indicates that the increased β-catenin in osteoblasts by tibial loading is attenuated in Δ130-136 and the Cx43(M1)-treated mice. These results establish a close functional relationship between Cx43 hemichannel-released PGE_2_ and decreased Sost, and thereby increased β-catenin expression in osteoblasts, ultimately leading to enhanced osteoblast activity and endosteal bone formation (**Figure 8H**).

There are possible limitations in this study. First, analysis of cortical bone changes at additional proximal or distal sites may provide a more comprehensive understanding of the role of Cx43 hemichannels in anabolic responses to mechanical loading, although a previous study has reported that cortical bone located at 25%, 37% and 50% of the tibia’s length had similar responses to tibial loading (Yang et al., 2017). Second, the monoclonal Cx43(M1) antibody blocks hemichannels not only in osteocytes, but also, possibly other cells, such as osteoblasts. However, in our study, Cx43(M1) was primarily detected in osteocytes, not in osteoblasts or other bone cells.

In summary, this study, for the first time, unveils the crucial role of osteocytic Cx43 hemichannels in mediating the anabolic function of mechanical loading on endosteal bone surfaces and trabecular bone. Cx43 hemichannels activated by mechanical stimulation release PGE_2_ from osteocytes, which suppresses Sost expression in osteocytes, and enhances osteoblast activity and bone formation on endosteal surfaces. These results suggest that osteocytic Cx43 hemichannels could be established as a *de novo* new therapeutic target, and activation of these channels may potentially aid in treating bone loss, in particular, in the elder population with the lost sensitivity to anabolic responses to mechanical stimulation (Lanyon and Skerry, 2001).

## Materials and Methods

### Mouse models

Two transgenic models expressing dominant-negative mutants of Cx43 in osteocytes, R76W and Δ130-136, were generated as previously described (Xu et al., 2015). The two transgenes were driven by a 10-kb DMP1 promoter and expressed predominantly in osteocytes. The WT and transgenic mice in C57BL/6J background were housed in a temperature-controlled room with a light/dark cycle of 12 hrs at the University of Texas Health Science Center at San Antonio (UTHSCSA) Institutional Lab Animal Research facility under specific pathogen-free conditions. Food and water were freely available. 15-week-old male WT and homozygous transgenic were sedated under isoflurane and euthanized by cervical dislocation. All animal protocols were performed following the National Institutes of Health guidelines for care and use of laboratory animals and approved by the UTHSCSA Institutional Animal Care and Use Committee (IACUC).

### Tibial mid-diaphyseal strain measurements and cyclic tibial loading

The relationship between applied compressive loading and bone tissue deformation of the left tibia was established for 15-week-old mice *in vivo* following a previously reported protocol (De Souza et al., 2005; Lynch et al., 2010). Briefly, a strain gauge (EA-06-015DJ-120, Vishay Measurements Group) was attached on the tibial diaphyseal medial mid-shaft of a euthanized mouse and load applied from 0 to 9.5 N at the ends of the left tibia using a loading machine (LM1, Bose). The strain gauge was connected to a bridge completion module (MR1-350-127, Vishay Measurements Group) and a conditioner/amplifier system. Mechanical load-induced strain was measured, and the compliance relationship between applied load and the resulting strain was determined for each left tibia (R^2^ > 0.99). Compared to WT mice, a higher compressive force was required to generate comparable periosteal strain in Δ130-136 mice (**Figure 1-figure supplement 1C**).

The cyclic axial compressive load was applied to the left tibia of each mouse using a custom loading device based on previous studies (De Souza et al., 2005; Lynch et al., 2010). Briefly, the left tibia of anesthetized mice was positioned into a custom-made apparatus (**Figure 1-figure supplement 1A**). The upper padded cup containing the knee was connected to the loading device (7528-10, Masterflex L/S, Vernon Hills, IL, USA), and the lower cup held the heel. The left tibia was held in place by a 0.5 N continuous static preload, loaded for 600 cycles (5 min) at 2-Hz frequency, with a sinusoidal waveform (**Figure 1-figure supplement 1B**). Compressive load was performed 5 days/week for 2 weeks to determine bone structural and anabolic response, or 5 consecutive days to assess PGE_2_ level. Based on the load-strain relationship, peak force was selected to generate peak periosteal strains of 1200 με at the cortical midshaft for WT, R76W, and Δ130-136 mice, respectively. This strain level has been previously shown to elicit an anabolic response at this region(Melville et al., 2015). The right tibia was used as a contralateral, non-loaded control.

### M1 antibody generation and treatment

A monoclonal Cx43(M1) antibody targeting the second extracellular loop domains of Cx43 was originally generated by Abmart (Tulsa, OK, USA) and described previously (Zhang et al., 2021). Briefly, mice were immunized with a Cx43 extracellular domain peptide, and after functional characterization of the hybridoma clones, genes that encode the antibody heavy and light chain variable region were cloned from the mouse hybridoma cell line HC1 by reverse transcription quantitative PCR (RT-qPCR), using a combination of a group of cloning PCR primers. The heavy and light chain constructs were co-transfected into human embryonic kidney freestyle 293 (HEK293F) cells, supernatants were harvested, and antibodies were purified by affinity chromatography using protein A resin.

The day before tibial loading, randomly allocated WT mice based on the body weight were intraperitoneally (IP) injected with 25 mg/kg Cx43(M1) or vehicle (phosphate-buffered saline (PBS), pH 7.4). A second dose was administered the day before the start of loading in the 2nd week. The dosage of antibody was based on our data with Cx43(M1) antibody (**Figure S5**) and a previous study using an anti-sclerostin (Sost) antibody (Spatz et al., 2013).

### *In vitro* dye uptake and gap junction coupling assays

The osteocyte-like MLO-Y4 cells were cultured in a-modified essential medium (a-MEM) with 2.5% fetal bovine serum (FBS) and 2.5% calf serum (CS) in a 5% CO_2_ incubator at 37℃. MLO-Y4 cells were grown at a low initial cell density on glass slides coated with type I collagen (rat tail collagen type I, Corning, Bedford, MA, USA, 0.15 mg/ml) to ensure that most of the cells were not physically in contact. The cells were preincubated with Cx43(E2) or Cx43(M1) (2 μg/ml) for 30 min and then subjected to FFSS at 4 dynes/cm^2^ for 15 min in the presence of 25 mM ethidium bromide (EtBr) in the recording media (HCO_3_^−^-free a-MEM medium buffered with 10 mM HEPES). These cells were then fixed with 1% paraformaldehyde (PFA) for 10 min. The intensity of EtBr fluorescence in cells was measured and quantified by NIH Image J software (NIH, USA). Primary osteocytes were microinjected using an Eppendorf micromanipulator InjectManNI 2 and Femtojet (Eppendorf) at 37°C with 10 mM Oregon green 488 BAPTA-AM (Mr: 1751 Da) as a cell tracker probe, and calcein red-orange AM (Mr: 789 Da) as a probe for detecting gap junction coupling. Images were captured using an inverted microscope equipped with a Lambda DG4 device (Sutter Instrument Co, Novato, CA, USA), a mercury arc lamp illumination, and a Nikon Eclipse microscope (Nikon, Tokyo, Japan) using a rhodamine filter. Loaded cells (dye donor) were “parachuted” over acceptor cells. The cells (acceptors and donors) were pre-incubated for 20 min with Cx43 antibodies before the parachuting assay. Donor cells were then incubated with acceptor cells for 90 min, the time duration sufficient to detect dye transfer.

### *In vivo* dye uptake assay

We developed an approach to assess hemichannel activity in osteocytes in the bone *in vivo* (Riquelme et al., 2021). Briefly, 20 mg/ml Evans blue (EB) dye dissolved in sterile saline solution (previously used to study hemichannel activity in muscle cells *in vivo* (Cea et al., 2013)) was injected into the mouse tail vein. For the vehicle or Cx43(M1) treated group, mice were IP injected with mouse IgG or Cx43 (M1) (25 mg/kg) 4 hrs before dye injection. After the dye injection, mice were kept in cages for 20 min, and the left tibias were then loaded for 10 min. Mice were sacrificed and perfused with PBS and 4% PFA 40 min after tibial loading. Tibias were isolated and fixed in 4% PFA for 2 days, decalcified in 10% ethylenediaminetetraacetic acid (EDTA) for 3 weeks, and then 12-μm-thick frozen sections were prepared. The cell nuclei were stained with 4’,6-diami-dino-2-phenylindole (DAPI). Images were captured using an optical microscope (BZ-X710, KEYENCE, Itasca, IL, USA) and EB fluorescence intensity in osteocytes was quantified by NIH Image J software (NIH, USA).

### PGE_2_ measurement and treatment

The level of PGE_2_ in the tibia bone was determined according to the manufacturer’s protocol (PGE_2_ ELISA kit, #514010, Cayman Chemical, Ann Arbor, MI, USA). Briefly, 4 hrs after the final round of five-day tibial loading, bone marrow-flushed tibias were isolated free of soft tissues, and bone shafts were prepared by removing proximal and distal ends of the bone. Bone tissue was homogenized in liquid nitrogen with a frozen mortar and pestle. The concentration of PGE_2_ was normalized by total protein concentration using a BCA assay (#23225, Thermo Scientific, Rockford, IL, USA).

PGE_2_ powder (#2296, Tocris Bioscience, Bristol, UK) was dissolved in 10% ethanol and stock prepared at the concentration of 0.15 mg/ml. Wild-type mice randomly allocated based on the body weight were injected with 1 mg/kg/day of PGE_2_ solution or 6.7μl/kg/day 10% ethanol (vehicle) for two weeks.

### Micro-computed tomography

Tibias were dissected and frozen in saline-soaked gauze at -20°C until scanning. Samples in PBS were imaged using a high-resolution micro-computed tomography (μCT) scanner (1172, SkyScan, Brüker microCT, Kontich, Belgium) with the following settings: 59 Kvp, 167 µA beam intensity, 0.5 mm aluminum filter, 800 ms exposure, 1024 x 1024 pixel matrix, and a 10 µm isotropic voxel dimension. The background noise was removed from the images by eliminating disconnected objects smaller than 4 pixels in size. Two bone volumes of interest (VOI) were selected in the metaphyseal and midshaft regions. The analyses were conducted excluding the fibula. In the proximal tibial metaphysis, the trabecular bone VOI was positioned 0.44 mm distal to the proximal growth plate and extended 0.65 mm in the distal direction, excluding the primary spongiosa. Grayscale values of 80-256 were set as the threshold for trabecular bone. For the cortical region, a 0.3 mm distance was centered at 50% tibial length (proximal to distal). Automated contouring was used to select the cortex. A threshold of 106-256 was applied to all of the cortical slices for analysis. The structural morphometric properties of cortical and trabecular regions were analyzed using the CT Analyser software (CTAn 1.18.8.0, Bruker Skyscan).

### Mechanical testing

Three-point bending tests were performed after μCT scanning. The tibia was thawed to room temperature before testing. Any remaining muscles and the fibula were carefully removed. The tibia was subjected to a three-point bending test along the medial-lateral direction in a micromechanical testing system (Mach-1 V500CST, Biomomentum, Laval, Canada). The span distance for the three-point bending test was 8 mm, and the loading pin was placed at the midpoint of the span. The test was performed in a displacement control mode at a constant rate of 0.01 mm/sec, and the data was collected at a 200-Hz sampling rate for all measurements. The accurate cross-sectional areas were determined from μCT and used to calculate mechanical properties (Jepsen et al., 2015).

### Histomorphometry, immunohistochemistry, and dynamic bone histomorphometry

Tibias were collected and fixed in 4% PFA for 2 days and decalcified using 10% EDTA (pH 7.5) for 21 days. These tibial samples were embedded in paraffin, and 5-mm-thick sections were collected and mounted onto glass slides. For static bone histomorphometry, tartrate resistant acid phosphatase (TRAP) and toluidine blue staining was used to determine the osteoclast activity (Xu et al., 2015) and osteoblast numbers, respectively. For immunohistochemistry, paraffin sections were rehydrated and antigen site was retrived with 10 mM citrate buffer (pH 6.0) at 60°C for 2 hrs (for sclerostin or β-catenin) or trypsin buffer (pH 7.8) at 37°C for 30 min (for COX-2), and then probed with an anti-sclerostin (AF1589, 1:400, R&D systems, Minneapolis, MN, USA), anti-COX-2 (12375-1-AP, 1:200, Proteintech, Rosemont, IL, USA) or an anti-β-catenin antibody (ab16051, 1:200, Abcam, Waltham, MA, USA) overnight at 4°C. The sections were probed with a biotin-labeled secondary antibody and ABC Reagent (VECTASTAIN, Burlingame, CA, USA). Staining was visualized with DAB Chromogen (SK-4100, Vector Laboratories, Burlingame, CA, USA). Hematoxylin was used as a counterstain. Images were captured using an optical microscope (BZ-X710, KEYENCE). Osteoclast surface (Oc.S), osteoclast number (N.Oc), osteoblast number (N.Ob), and bone surface (BS) on the tibial midshaft cortical bone along the endosteal surface were counted and positive cells quantified using ImageJ software (NIH, USA).

Mice were IP injected with calcein (C0875, Sigma-Aldrich, St. Louis, MO, USA) at 20 mg/kg of body weight 1 day before tibial loading, and followed by alizarin red injection (A5533, Sigma-Aldrich, St. Louis, MO, USA) at 30 mg/kg of body weight 3 days before euthanization. Tibias were dissected, fixed in 70% ethanol, and then embedded in methylmethacrylate for 10-μm thick longitudinal plastic sections. Two-color fluorescent images were obtained using a fluorescence microscope (BZ-X710, KEYENCE). Single label was defined as any bone surface with green, red, or yellow (no separation between green and red). The distance between green and red fluorescence signals was measured along the bone surface (Grimston et al., 2012). The following parameters were quantified at tibial midshaft using NIH ImageJ software (NIH, USA): total perimeter (BS); single label perimeter (sLS); double label perimeter (dLS), and double-label area (dL.Ar). The following values were then calculated: mineralizing surface [MS/BS = (sLS/2 + dLS)/BS], mineral apposition rate [MAR = dL.Ar/dLS/12], and bone formation rate (BFR/BS = MAR × MS/BS).

### RNA extraction and RT-qPCR

The long tibial bone was isolated free of soft tissues, and bone marrow was removed by flushing with RNase-free PBS after two weeks tibial loading. The bone shaft was prepared by removing the proximal and distal ends of the bone and pulverizing it using a frozen mortar and pestling in liquid nitrogen. Total RNA was isolated by using TRIzol (Molecular Research Center, Cincinnati, OH, USA) according to the manufacturer’s protocol. cDNA was synthesized by a high-capacity cDNA reverse transcription kit (#4388950, Applied Biosystems, Carlsbad, CA, USA). mRNA level was analyzed by real-time RT-qPCR using an ABI 7900 PCR device (Applied Biosystems, Bedford, MA, USA) and SYBR Green (#1725124, Bio-Rad Laboratories, Hercules, CA, USA) with a two-step protocol (94°C for 10 sec, and 65°C for 30 sec for 40 cycles). The relative gene expression in each loaded tibia was represented by normalizing to GAPDH and then normalized to the control tibia (2^-ΔΔCt^) (Kenneth J. Livak and Schmittgen, 2001). The primers for *Sost*, *COX-2, Runx2, Bgalp, β-catenin,* and *Dmp1* are provided in Table S1.

### Statistical analysis

Data collection and analysis were conducted in a blind manner. Statistical analysis was performed using IBM SPSS Statistics 24 (SPSS Inc., Chicago, IL, USA) and graphed with GraphPad Prism 7 (GraphPad Software; La Jolla, CA, USA). For *in vitro* studies, each experiment had three technical replicates and was repeated at least three times. Normal distribution of the data was evaluated by the Shapiro-Wilk test, and homogeneity of variance was assessed by the Levene test. The paired t-test was used for comparisons of the loaded and contralateral tibias within the same group. One-way ANOVA with Tukey test was used for multiple group comparisons. Student unpaired t-test was used to compare between vehicle and Cx43 (M1)-treated groups. All data are presented as means ± SD. P<0.05 was considered significant.

## Acknowledgements

We thank Dr. Eduardo Cardenas and Dr. Francisca Acosta at UTHSCSA for critical reading and editing of the paper.

## Funding

This work was supported by the National Institutes of Health (NIH) Grants: 5RO1 AR072020 (to J.X.J.), and Welch Foundation grant: AQ-1507 (to J.X.J.). We also thank the support of the UTHSCSA CMMI and the UTHSCSA Optical Imaging Facility supported by the Cancer Therapy and Research Center through NIH–National Cancer Institute P30 award CA054174 and Texas State funds.

## Conflicts of interest/Competing interests

The authors declare no conflicts of interest with the contents of this article.

## Authors’ contributions

D.Z., G.S. and J.X.J. designed the study; D.Z., M.A.R., T.G., L.W., C.T., H.X., G.S. performed experiments and analyzed the data; D.Z., and J.X.J. wrote the manuscript, which was reviewed, commented and approved by all authors.

**Figure 1-Source Data 1.** Trabecular micro-CT data of transgenic and wild-type mice.

**Figure 1-figure supplement 1-source data 1.** Raw data of compliances for Figure 1-figure supplement 1C.

**Figure1-figure supplement 2-source data 1.** Raw data of dye uptake for Figure 1-figure supplement 2B.

**Figure1-figure supplement 3-source data 1.** Raw data of body weight for Figure1-figure supplement 3A.

**Figure 2-Source Data 1**. Cortical micro-CT data of transgenic and wild-type mice.

**Figure 3-Source Data 1.** Raw data of periosteal and endosteal bone formation of transgenic and wild-type mice.

**Figure 4-Source Data 1.** Raw data of PGE_2_ level for Figure 4A.

**Figure 4-Source Data 2.** Raw data of immumohistochemical, TRAP and toluidine blue staining of transgenic and wild-type mice.

**Figure 4-Source Data 3.** Raw data of RT-qPCR of transgenic and wild-type mice.

**Figure 4-figure supplement 1-source data 1.** Raw data of COX-2 and Sost quantification for Figure 4-figure supplement 1A, B.

**Figure 5-Source Data 1.** Micro-CT data of vehicle and Cx43(M1)-treated mice.

**Figure 5-Source Data 2.** Three-point bending data of vehicle and Cx43(M1)-treated mice.

**Figure 5-figure supplement 1-source data 1** Raw data of dye uptake for Figure 5-figure supplement 1B and F.

**Figure 5-figure supplement 2-source data 1.** Raw data of body weight for Figure 5-figure supplement 2A.

**Figure 6-Source Data 1.** Raw data of periosteal and endosteal bone formation of vehicle and Cx43(M1)-treated mice.

**Figure 7-Source Data 1.** Raw data of PGE_2_ level for Figure 7A.

**Figure 7–Source Data 2.** Raw data of immumohistochemical, TRAP and toluidine blue staining of vehicle and Cx43(M1)-treated mice.

**Figure 7-Source Data 3.** Raw data of RT-qPCR of vehicle and Cx43(M1)-treated mice.

**Figure 7-figure supplement 1-source data 1.** Raw data of COX-2 Sost and β-catenin quantification for Figure 7-figure supplement 1A-C.

**Figure 8-Source Data 1.** Cortical micro-CT data of vehicle and Cx43(M1)-treated mice treated with 1 mg/kg/day PGE2 or vehicle control.

**Figure 8-figure supplement 1-source data 1.** Raw data of body weight for Figure8-figure supplement 1A.

**Figure 8-figure supplement 2-source data 1.** Trabecular micro-CT data of vehicle and Cx43(M1)-treated mice treated with 1 mg/kg/day PGE2 or vehicle control.

---

**Figure 1-figure supplement 1.**
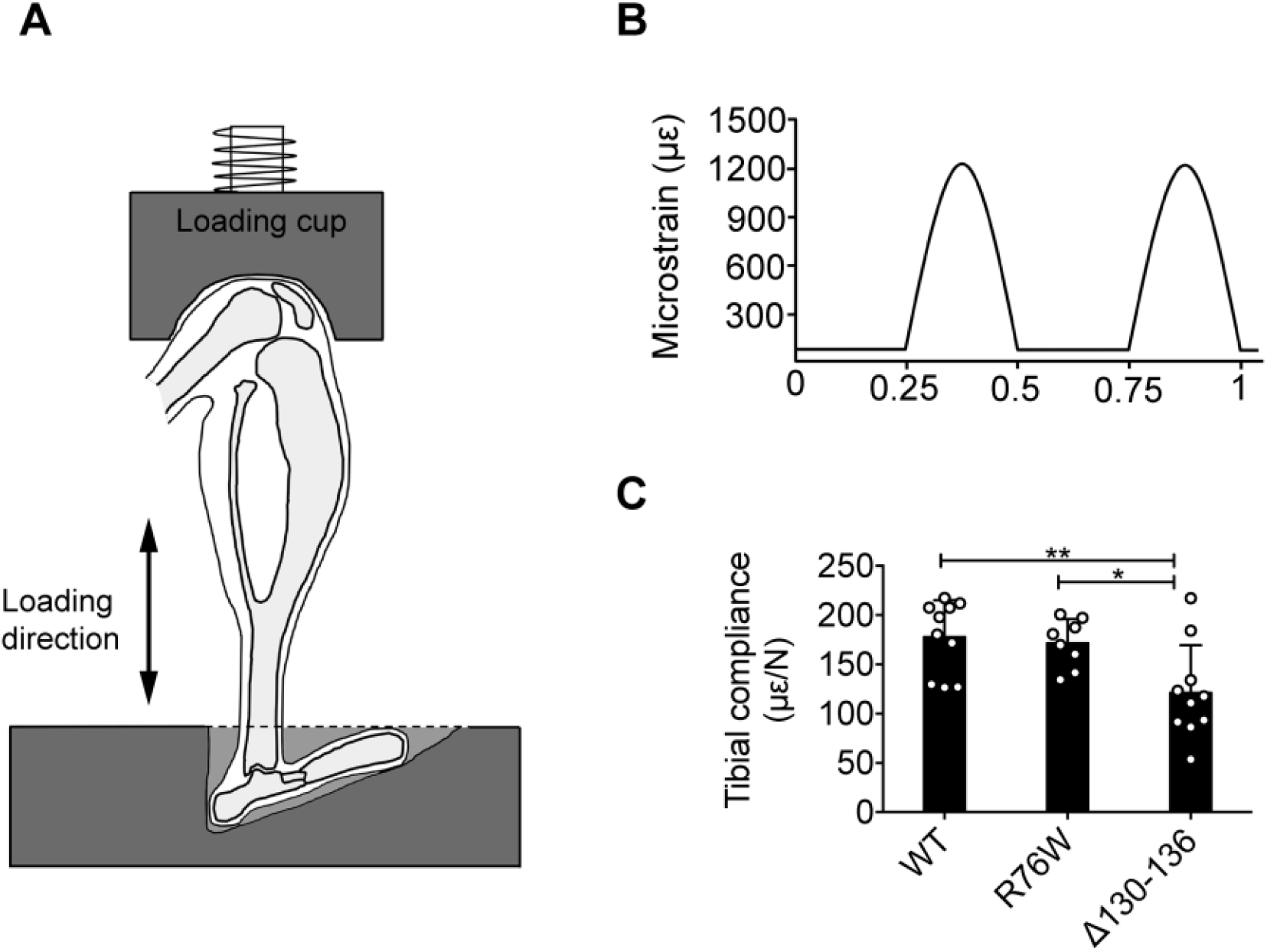
Experimental setup for *in vivo* axial loading. **(A)** Diagram of the left tibia positioned at the loading device and the direction of loading. **(B)** Schematic graph of 1 s of the daily 5 min loading signal. Approximately 1200 microstrain was detected on the medial mid-shaft surface of the tibia. **(C)** The average compliance of the relationship between applied load and resulting strain on the medial mid-shaft of WT, R76W and Δ130-136 mice. n=8-10/group. Data are expressed as mean ± SD. *, P<0.05; **, P<0.01. Statistical analysis was performed using one-way ANOVA with Tukey test among groups with different genotypes.

**Figure 1-figure supplement 2.**
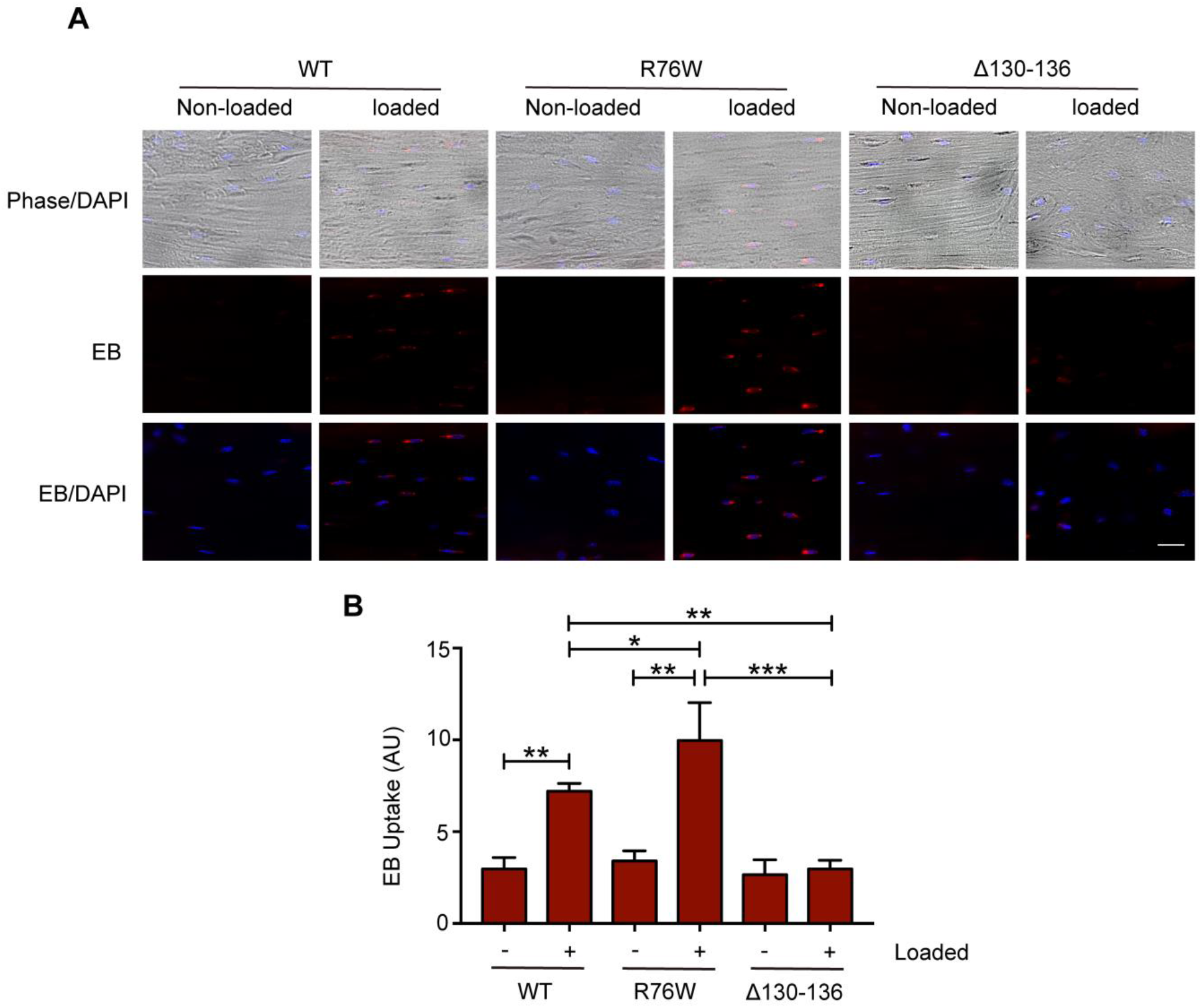
Hemichannel opening is Inhibited in Δ130-136 mice. (**A**) Representative images of Evans blue (EB) dye uptake in control and loaded tibial bone in WT, R76W and Δ130-136 mice. Scale bar, 60 μm. **(B)** Quantitative analysis of Evans blue (EB) dye uptake. n=3/group. Data are represented as mean ± SD. *, P<0.05; **, P<0.01; ***, P<0.001. Statistical analysis was performed using one-way ANOVA with Tukey test among different groups.

**Figure 1-figure supplement 3.**
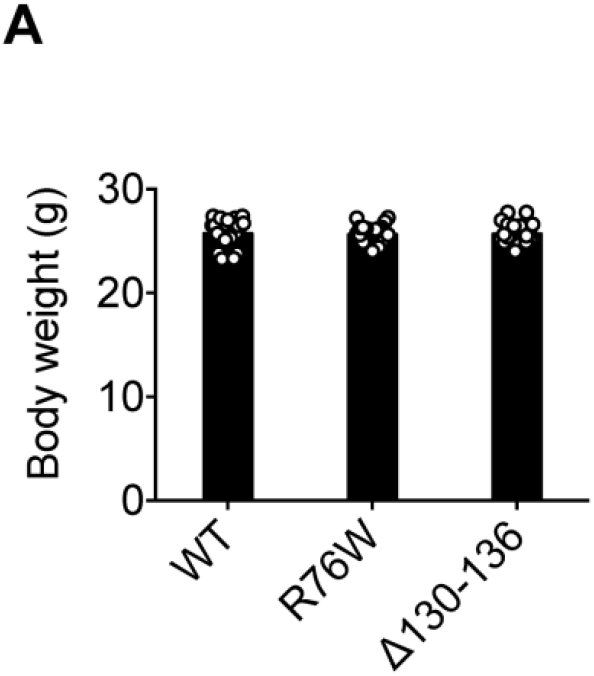
Body weights of transgenic mice. **(A)** The body weights of WT, R76W and Δ130-136 mice at the beginning of mechanical loading. n=22/group. Data are expressed as mean ± SD. Statistical analysis was performed using one-way ANOVA with Tukey test among different genotypes.

**Figure 4-figure supplement 1.**
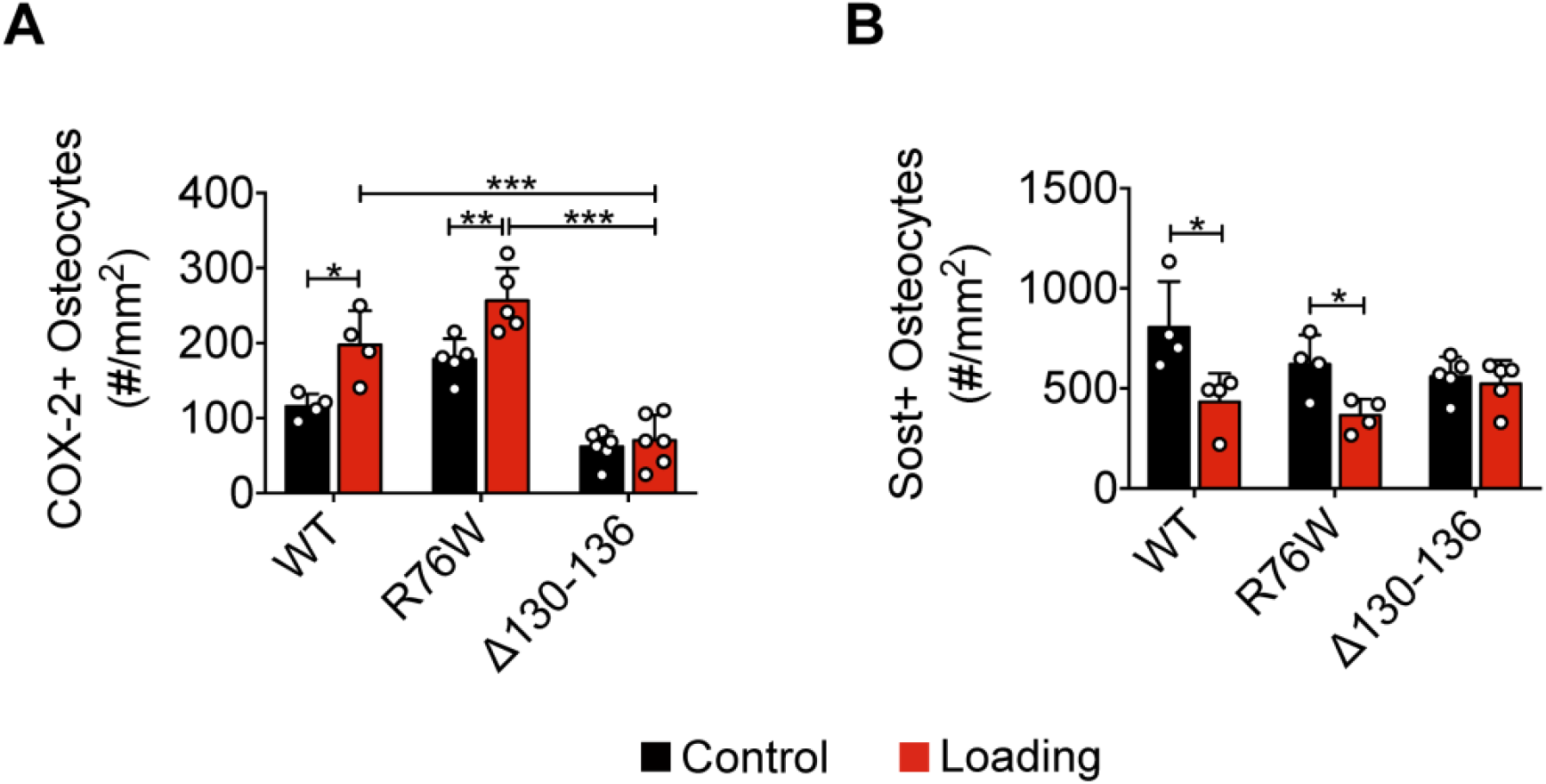
Bone marker protein expression in WT, R76W and Δ130-136 mice. **(A)** Quantitative analysis of the COX-2-positive osteocytes per bone area in tibial midshaft cortical bone after 2 weeks of mechanical loading in WT, R76W and Δ130-136 mice. n=4-6/group. (**B**) Quantitative analysis of the Sost -positive osteocytes per bone area in tibial midshaft cortical bone after 2 weeks of mechanical loading in WT, R76W and Δ130-136 mice. n=4-6/group. Data are expressed as mean ± SD. *, P<0.05; **, P<0.01; ***, P<0.001. Statistical analysis was performed using paired t test for loaded and contralateral tibias, or one-way ANOVA with Tukey test for loaded tibias among groups with different genotypes.

**Figure 5-figure supplement 1.**
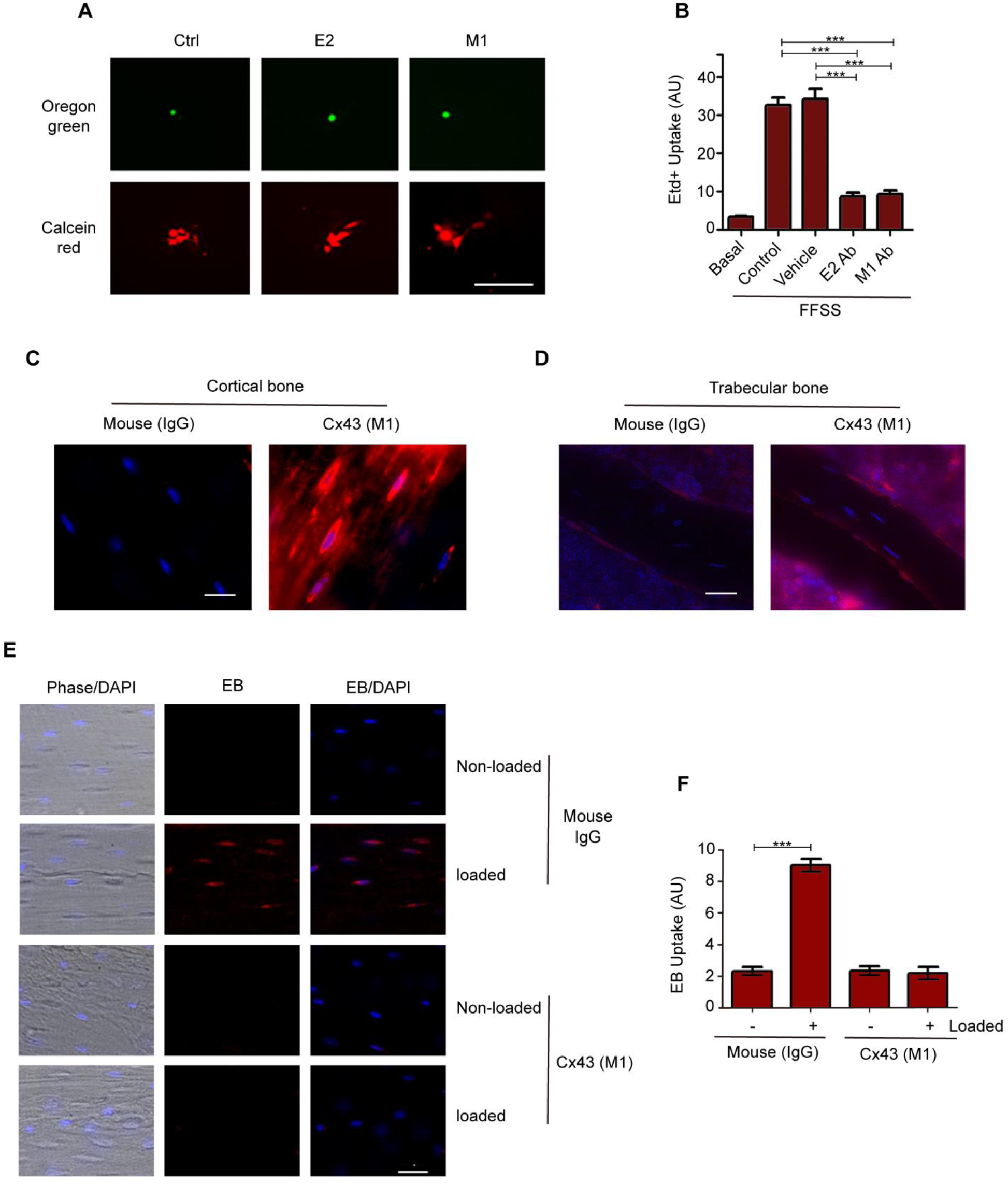
Monoclonal antibody of Cx43 inhibits hemichannel opening induced by mechanical stress *in vitro and in vivo*. **(A)** Parachuting dye coupling assay was conducted to determine gap junction coupling in MLO-Y4 cells loaded with Oregon green 488 BAPTA-AM (Mr: 1751 Da) as a cell tracker probe and calcein red-orange AM (Mr: 789 Da). Scale bar, 100 μm. **(B)** MLO-Y4 cells were preincubated with Cx43(E2), Cx43(M1), PBS (vehicle) or rhodamine-conjugated anti-mouse IgG (control) and then subjected to fluid flow shear stress (FFSS) (8 dynes/cm^2^) for 10 min and followed by ethidium bromide (Etd^+^) dye uptake assay. n=4/group. **(C and D)** Representative images of Cx43(M1) detected with rhodamine-conjugated anti-mouse IgG in tibial midshaft cortical bone and trabecular bone. Bar, 50 μm. **(E)** Representative images of Evans blue (EB) dye uptake in control and tibial loaded bone in the absence or presence of Cx43(M1) antibody. Scale bar, 40 μm. **(F)** Quantitative analysis of Evans blue (EB) dye uptake. n=3/group. Data are expressed as mean ± SD. *, P<0.05; **, P<0.01; ***, P<0.001. Statistical analysis was performed using one-way ANOVA with Tukey test among different groups.

**Figure 5-figure supplement 2.**
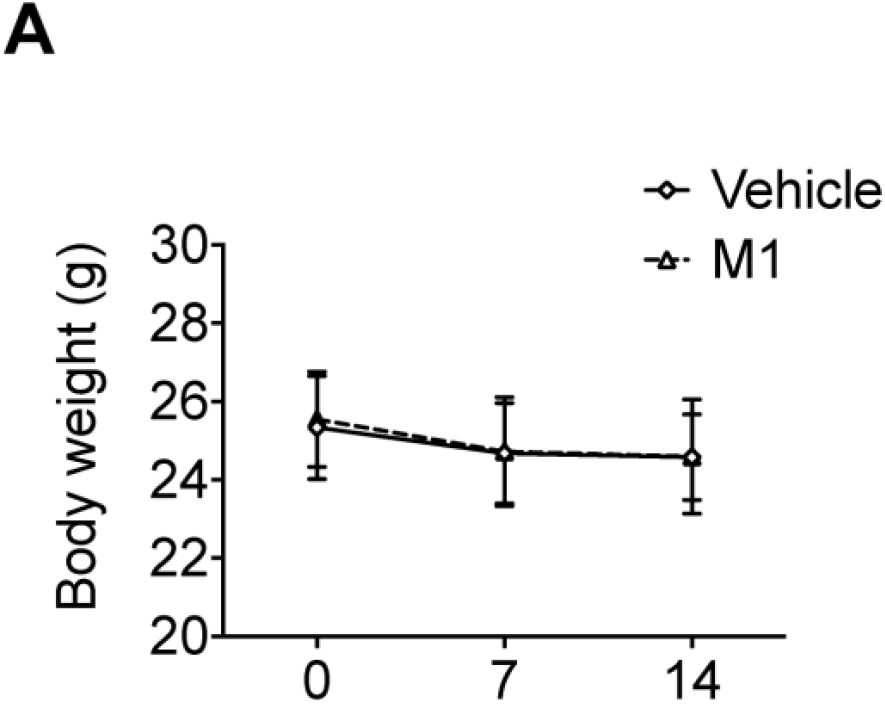
Body weights of mice during 2 weeks of tibial loading. **(A)** Weekly body weights of vehicle and Cx43(M1)-treated groups. n=18/group. Data are expressed as mean ± SD. Statistical analysis was performed using one-way ANOVA with Tukey test among different genotypes.

**Figure 7-figure supplement 1.**
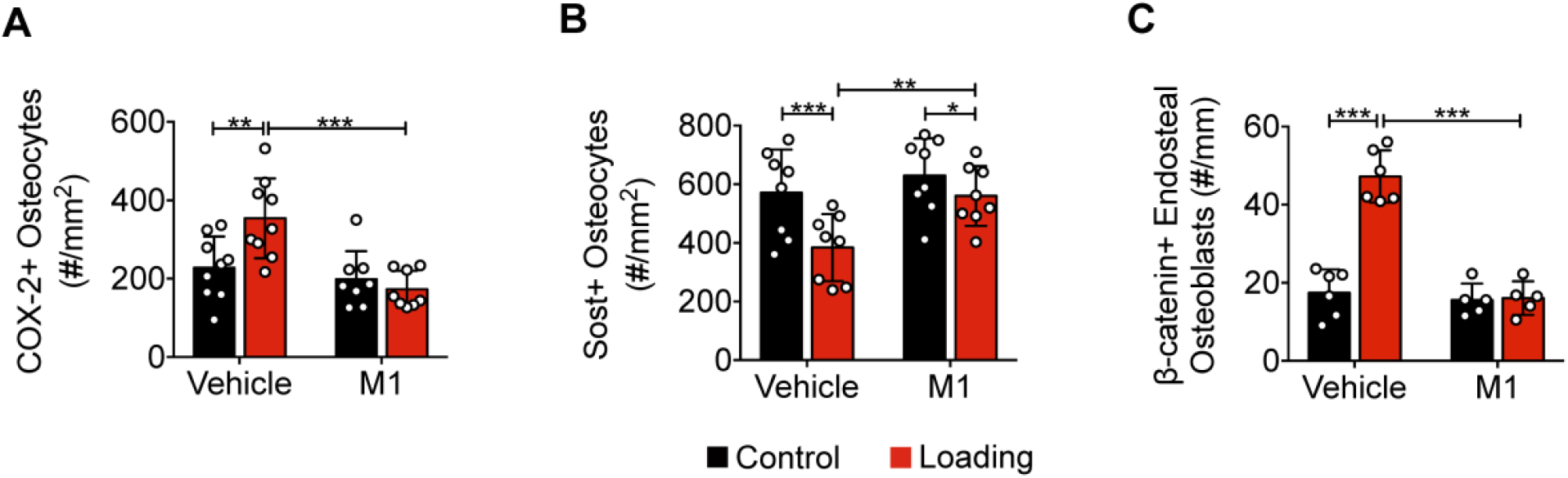
Bone marker protein expression in vehicle- and Cx43(M1)-treated mice. **(A, B)** Quantitative analysis of the COX-2-positive or Sost-positive osteocytes per bone area in tibial midshaft cortical bone after 2 weeks of loading in vehicle and Cx43(M1)-treated mice. n=8-9/group. **(C)** Quantitative analysis of the β-catenin-positive osteoblasts per bone perimeter in tibial midshaft cortical bone after 2 weeks of mechanical loading in vehicle and Cx43(M1)-treated mice. n=4-6/group. Data are expressed as mean ± SD. *, P<0.05; **, P<0.01; ***, P<0.001. Statistical analysis was performed using paired t test for loaded and contralateral tibias, unpaired t test for loaded tibias between vehicle- and Cx43(M1)-treated groups.

**Figure 8-figure supplement 1.**
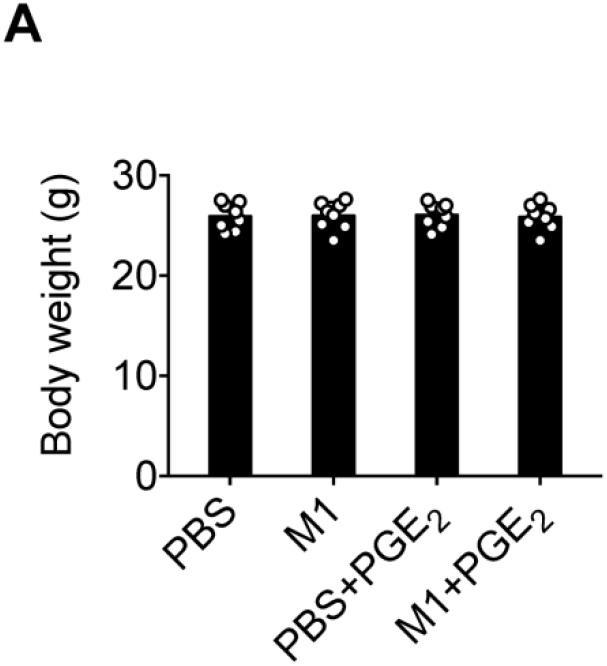
Body weights of mice of vehicle- and Cx43(M1)-treated mice treated with 1mg/kg/day PGE2 or vehicle control. **(A)** The body weights of vehicle- and Cx43(M1)-treated mice treated with 1mg/kg/day PGE_2_ or vehicle control at the beginning of mechanical loading. n=8/group. Data are expressed as mean ± SD. Statistical analysis was performed using one-way ANOVA with Tukey test among different genotypes.

**Figure 8-figure supplement 2.**
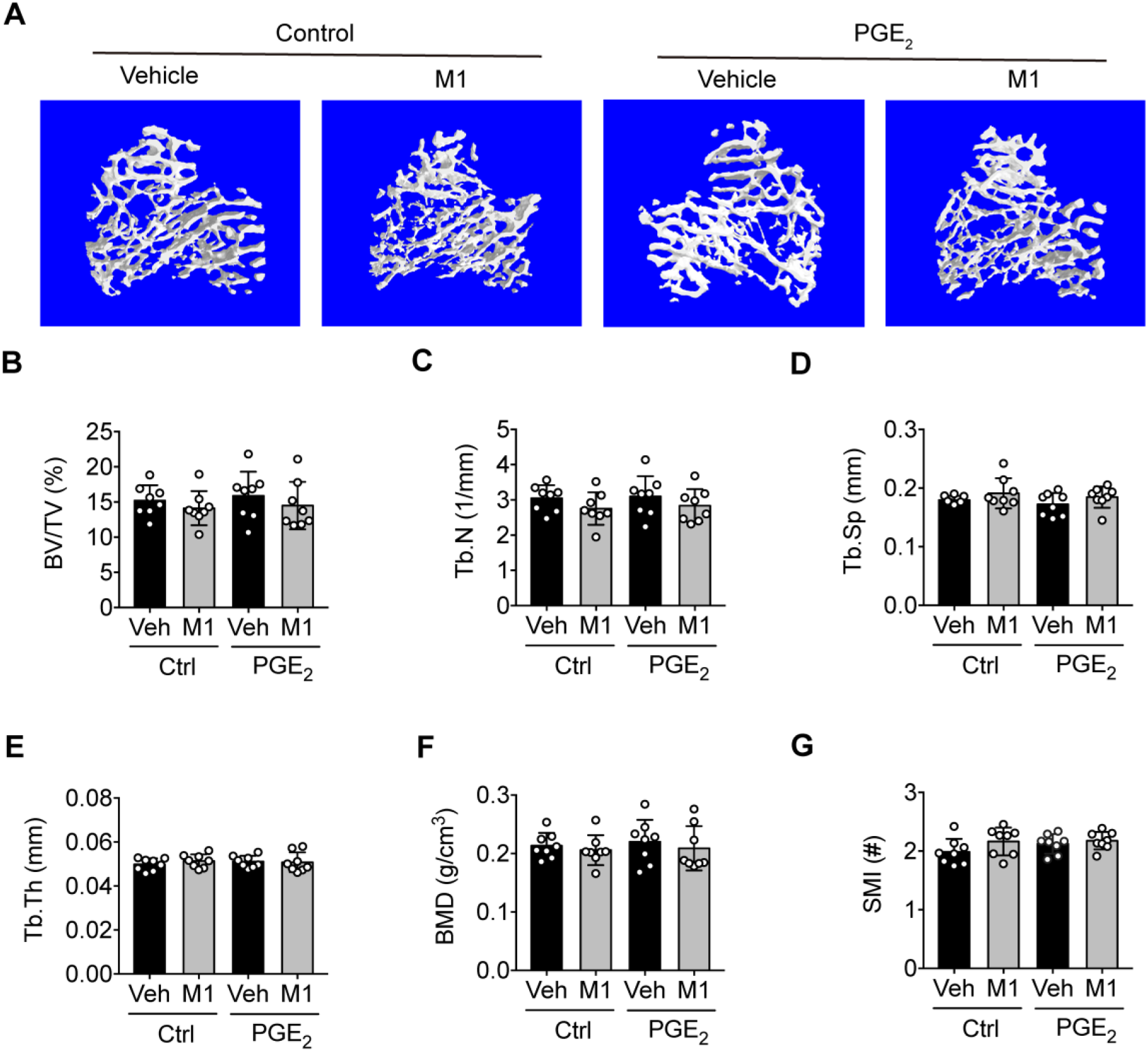
PGE_2_ does not exert additional trabecular osteogenic response to mechanical loading. **(A)** Representative 3D models of metaphyseal trabecular bone in vehicle and Cx43(M1)-treated mice treated with 1 mg/kg/day PGE_2_ or vehicle control. **(B-G)** μCT was used to assess tibial midshaft cortical bone; **(B)** bone volume fraction, **(C)** trabecular number, **(D)** trabecular separation, **(E)** trabecular thickness, **(F)** bone mineral density and **(G)** structure model index. n=8/group. Data are expressed as mean ± SD. *, P<0.05; **, P<0.01; ***, P<0.001. Statistical analysis was performed using one-way ANOVA with Tukey test among different groups.

## Notes

### Competing Interest Statement

The authors have declared no competing interest.

